# Distinct mechanisms for sebaceous gland self-renewal and regeneration provide durability in response to injury

**DOI:** 10.1101/2023.05.05.539454

**Authors:** Natalia A. Veniaminova, Yunlong Jia, Adrien M. Hartigan, Thomas J. Huyge, Shih-Ying Tsai, Marina Grachtchouk, Seitaro Nakagawa, Andrzej A. Dlugosz, Scott X. Atwood, Sunny Y. Wong

## Abstract

Sebaceous glands (SGs) release oils that protect our skin, but how these glands respond to injury has not been previously examined. Here, we report that SGs are largely self-renewed by dedicated stem cell pools during homeostasis. Using targeted single cell RNA-sequencing, we uncovered both direct and indirect paths by which these resident SG progenitors ordinarily differentiate into sebocytes, including transit through a PPARγ+Krt5+ transitional cell state. Upon skin injury, however, SG progenitors depart their niche, reepithelialize the wound, and are replaced by hair follicle-derived stem cells. Furthermore, following targeted genetic ablation of >99% of SGs from dorsal skin, these glands unexpectedly regenerate within weeks. This regenerative process is mediated by alternative stem cells originating from the hair follicle bulge, is dependent upon FGFR signaling, and can be accelerated by inducing hair growth. Altogether, our studies demonstrate that stem cell plasticity promotes SG durability following injury.

## INTRODUCTION

Our skin is coated with a complex mixture of oils that serves critical roles in modulating water retention, body temperature and the microbiome. These oily secretions, known as sebum, originate from sebaceous glands and constitute up to 90% of the total surface lipids in the skin (Niemann and Horsley, 2012; Zouboulis et al., 2022). Over-production of sebum by sebaceous glands can lead to “oily skin,” whereas hyposecretion of sebum is often associated with dry skin and eczematous dermatoses (Lovászi et al., 2017; Shi et al., 2015). Since perturbations in sebum are notably linked to common cutaneous disorders such as acne, seborrheic dermatitis and enlarged facial pores, sebaceous glands must be exquisitely regulated in order to maintain healthy skin function and cosmetic appearance (Lee et al., 2016; Smith and Thiboutot, 2008).

Sebaceous glands (SGs) are epithelial appendages typically associated with hair follicles. These acinar structures are comprised of terminally differentiated sebocytes ensheathed by a peripheral layer of undifferentiated basal progenitor cells (Cottle et al., 2013; Hinde et al., 2013). During maturation, sebocytes enlarge, accumulate lipids and degrade their organelles in a specialized form of cell death known as holocrine secretion (Fischer et al., 2017). This process culminates with sebocytes releasing their lipid contents through the sebaceous duct into the hair follicle infundibulum, which provides a passageway for sebum to exit the follicle and enter the skin surface (Schneider and Paus, 2014).

Since SGs are hormonally-regulated, their activity varies at different stages of life (Zouboulis and Boschnakow, 2001). Nonetheless, the constant turnover of sebocytes necessitates that these cells be continually replenished, a process that typically takes 1-2 weeks in mice and 2-4 weeks in humans (Epstein and Epstein, 1966; Jung et al., 2015; Plewig and Christophers, 1974; Weinstein, 1974). This renewal process is made possible by stem cells, although the niche in which these cells reside has not been decisively established. Whereas some studies have noted that hair follicle stem cells can enter and renew the gland (Han et al., 2023; Panteleyev et al., 2000; Petersson et al., 2011), other reports have indicated that SGs harbor their own dedicated stem cell pools (Andersen et al., 2019; Füllgrabe et al., 2015; Ghazizadeh and Taichman, 2001; Veniaminova et al., 2019). In addition, it remains controversial whether all basal progenitors that line the SG periphery contribute equally to sebocyte formation. Finally, whether these cellular processes become altered after injury has not been explored.

Technical challenges have posed a major hindrance to answering these questions. Because SGs are lobular structures that exhibit cellular heterogeneity along multiple axes—including proximal-distal, as well as proximity to the sebaceous duct—the spatial and molecular relationships of sebocytes at different stages of differentiation have been difficult to resolve. In addition, the lack of Cre drivers that specifically and efficiently target SGs complicates genetic fate mapping studies. Indeed, current tools for performing lineage tracing on SGs rely on mouse Cre lines that also target the hair follicle (Füllgrabe et al., 2015; Kretzschmar et al., 2014; Page et al., 2013; Petersson et al., 2011). Sebocytes are also challenging to isolate due to their complex cellular properties. Consequently, these cells typically constitute a minor sub-population in single cell RNA-sequencing (scRNA-seq) studies of skin, precluding the ability to perform deeper analyses (Cheng et al., 2018; Joost et al., 2020; Joost et al., 2016). Finally, studies on SG function have historically relied on mouse mutants such as *Asebia*, which possesses impaired SGs due to a germline mutation in *stearoyl-Coenzyme A desaturase-1* (*Scd1*) (Sundberg et al., 2000; Zheng et al., 1999).

Recent reports have suggested that SGs are adaptable structures that respond to local and systemic cues, are affected by the hair cycle and immune factors, and appear to be lost in diseases such as cicatricial alopecia and psoriasis (Choa et al., 2021; Karnik et al., 2009; Kobayashi et al., 2019; Reichenbach et al., 2018; Rittié et al., 2016; Stenn et al., 1999; Zhang et al., 2021). Here, we overcome many of the technical challenges for studying SGs, and perform highly targeted genetic fate mapping, scRNA-seq and ablation studies to explore how distinct stem cell populations maintain the gland and confer resiliency in response to injury.

## RESULTS

### Establishing SG landmarks

Keratins are by far the most abundant proteins in the skin, and the expression patterns of the 54 keratin family members subdivide keratinocytes by niche, function and differentiation status (Pan et al., 2013). We previously reported that basal progenitors at the SG periphery express keratins (K) 5 and 14, which form prototypic heterodimers in multiple mouse skin stem cell compartments (**Figure 1A**) (Veniaminova et al., 2019). In differentiated sebocytes, however, K14 levels remain high whereas expression of its typical binding partner K5 is reduced (**Figure 1A**). In its place, a different keratin, K79, becomes elevated in sebocytes and heterodimerizes with K14 (**Figure 1B**) (Veniaminova et al., 2019). Therefore, SG progenitors undergo a K14:K5 → K14:K79 keratin switch when they become sebocytes. Whether other keratins display similar shifts in the SG remains unclear and will be examined below.

**Figure 1.**
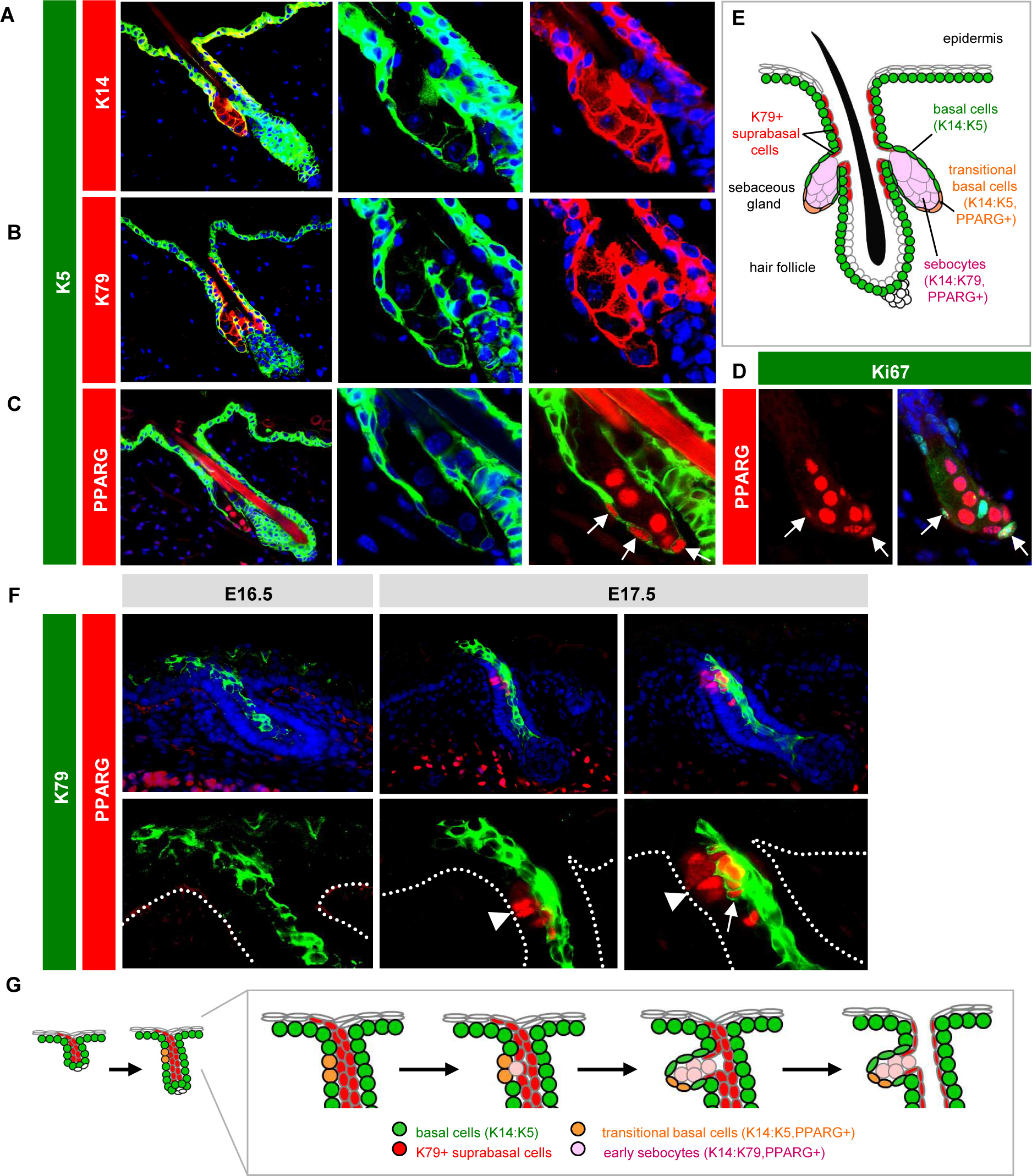
Establishing SG landmarks. **A.** Co-localization of K5 (green) with K14 (red) in peripheral SG basal cells, but not in sebocytes. Middle and right panels are magnified single-channel views. **B.** Lack of co-localization of K5 with K79 (red) in sebocytes. **C.** Co-localization of K5 with PPARγ (red) in transitional basal cells of the lower SG (arrows). **D.** Co-localization of PPARγ (red) with Ki67 (green) in a subset of peripheral basal cells (arrows) in the SG. **E.** Schematic of telogen hair follicle. Note that the infundibulum and sebaceous ducts are continuously lined by differentiated K79+ cells (red). **F.** Localization of K79 (green) and PPARγ (red) in the developing hair follicle during embryonic (E) days 16.5-17.5. Middle panels, follicle with basal PPARγ+ cells (arrowhead), but minimal co-localization with K79. Right panels, follicle with early sebocytes identified by the unique co-localization of PPARγ and K79 (arrow). Dotted lines delineate the basal layer of the epidermis and hair follicle. Bottom panels are magnified views, with DAPI omitted for clarity. **G.** Schematic of SG specification. PPARγ+ basal cells (orange) initially emerge at E16.5-17.5 and give rise to early sebocytes (pink) adjacent to the K79+ cell column (red). Subsequent remodeling leads to the opening of the sebaceous duct and hair canal (Mesler et al., 2020). One of two SG lobes is depicted. The second lobe may be specified later, or may arise when the initial SG compartment splits into two, as has been proposed (Frances and Niemann, 2012).

Notably, we also observed that peroxisome proliferator-activated receptor gamma (PPARγ), a master regulator of lipid metabolism and SG differentiation (Ahmadian et al., 2013; Ramot et al., 2015), is initially expressed by K5+ basal SG progenitors located at the lower (proximal) region of the gland (**Figure 1C**). A subset of basal PPARγ+ cells are also proliferative (**Figure 1D**). Because these cells express a unique combination of both progenitor (K5) and differentiation (PPARγ) markers, this suggests that these K5+PPARγ+ cells may behave as transitional basal cells poised to differentiate into sebocytes, a concept we revisit later (**Figure 1E**).

To determine whether these expression patterns are recapitulated during initial SG development, we examined mouse embryonic skin after hair follicle initiation but before SG formation. We have previously shown that nascent hair buds generate and extend columns of K79+ differentiated cells out into the epidermis, which subsequently undergo remodeling to form hair follicle openings (Mesler et al., 2020; Veniaminova et al., 2013). In embryonic (E) day 16.5 skin, we observed K79+ columns in developing hair buds, as expected, but no PPARγ expression (**Figure 1F**). By E17.5, however, we noticed basal PPARγ expression, reminiscent of the K5+PPARγ+ transitional basal cells seen in adult follicles (**Figure 1F**). Furthermore, we observed early sebocytes, identified by the unique co-expression of K79 and PPARγ, located immediately adjacent to the basal layer and alongside K79+ columns (**Figure 1F**). These findings are consistent with previous studies indicating that basal cells undergo asymmetric cell divisions to form sebocytes (Feldman et al., 2019; Frances and Niemann, 2012), and suggest a model for how the SG compartment becomes connected to the developing sebaceous duct and future hair follicle infundibulum—domains that are also unified by their shared expression of K79 (**Figure 1G**). In total, these observations establish a set of landmarks for evaluating SGs during homeostasis and injury.

### SG dynamics during skin homeostasis and injury

Given our observation that PPARγ is initially expressed in the SG basal layer, we next attempted to trace the fate of *Pparg*-expressing cells and their progeny. For this, we acquired AdipoTrak mice, in which a tetracycline-regulated transactivator (tTA) is expressed under the control of the endogenous *Pparg* promoter (**Figure 2A**) (Tang et al., 2008). When coupled with a tetracycline-responsive element (TRE)-driven Cre recombinase and a Cre-inducible YFP reporter allele (PPARγ;YFP mice), these genetic elements enable PPARγ+ cells and their descendants to become permanently labeled. However, in the presence of doxycycline (doxy), tTA cannot activate Cre expression, providing temporal control over this system.

**Figure 2.**
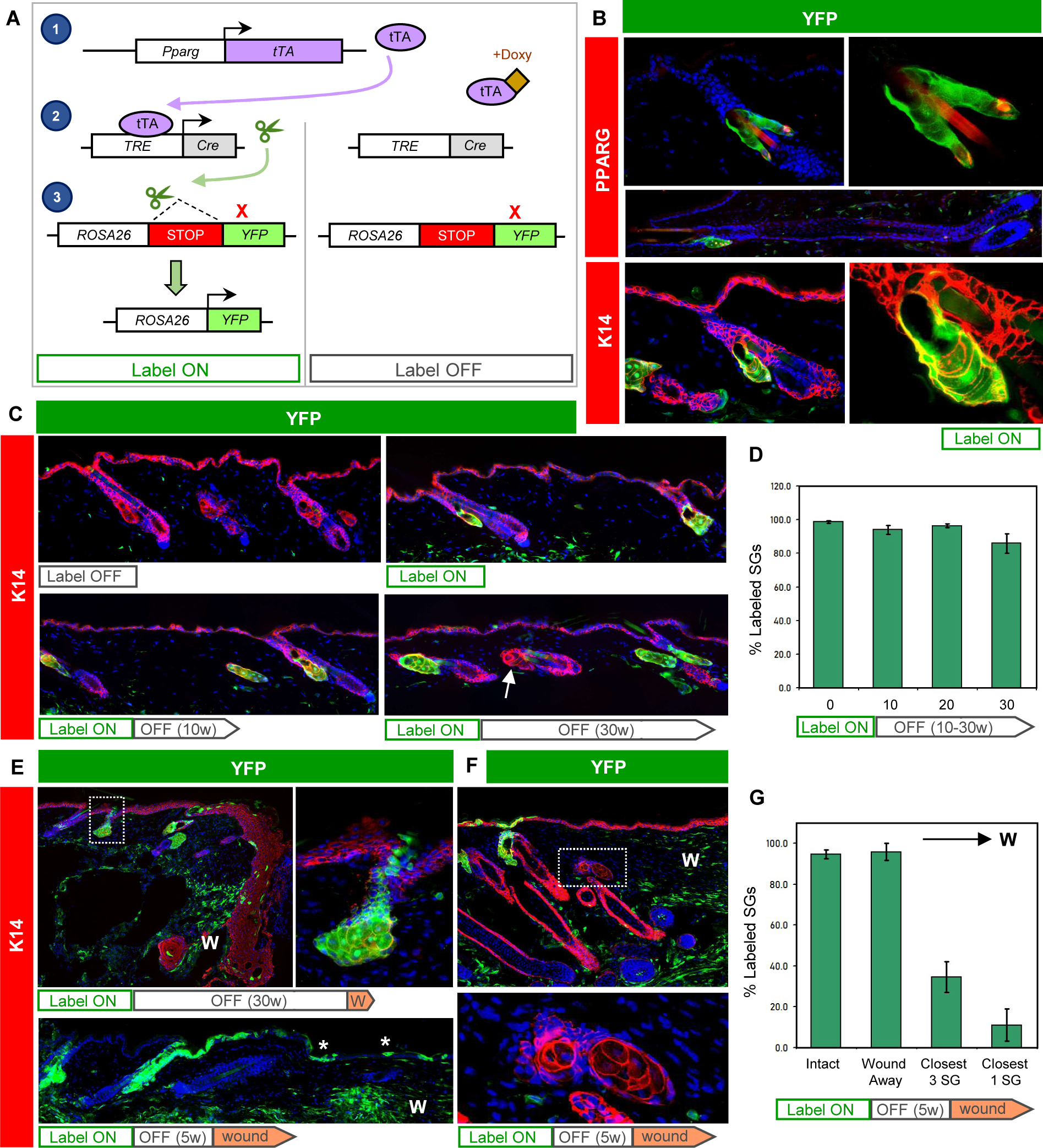
Tracing the SG during homeostasis and after wounding. **A.** Schematic for tracing PPARγ+ cells. Left, in the absence of doxycyline (doxy), *Pparg* promoter-driven tTA induces Cre expression, causing genomic recombination that activates YFP expression. Right, doxy suppresses tTA activity. **B.** Immunohistochemical localization of YFP (green) and PPARγ (red, top) or K14 (red, bottom) in label-on PPARγ;YFP mice. Basal SG cells, sebocytes and sebaceous ducts express YFP, but other hair follicle epithelia do not, in both telogen or anagen (middle). Right panels are magnified views of the left panels, with DAPI omitted. **C.** Top panels, 8 week old skin from label-off (left) or label-on (right) PPARγ;YFP mice. Bottom panels, skin from mice treated for the first time with doxy starting at 8 weeks of age, for 10-30 continuous weeks (label on→off). Arrow, unlabeled SG. **D.** Quantitation of labeled SGs, following 0-30 weeks of continuous doxy treatment. **E.** Wounded skin from a label on→off PPARγ;YFP mouse, examined 1 week (top) or 8 weeks (bottom) after injury. Top right panel is a magnified view of the boxed area showing labeled cells that have departed the SG and entered the epidermis. Asterisk, SG-derived YFP+ cells maintained long-term in the healed epithelium. **F.** Wounded skin from a label on→off PPARγ;YFP mouse, examined 3 weeks after injury. Bottom panel is a magnified view of the boxed area showing unlabeled, wound-proximal SGs. **G.** Quantitation of SG labeling, as a function of distance from the wound site. The closest SG cluster to the wound site is designated “closest 1,” and the closest 3 SG clusters are designated “closest 3.” W, wound site. w, weeks.

We began by analyzing 8 week old PPARγ;YFP mice without doxy exposure (label-on), and observed that >98% of all SGs were completely labeled (**Figure 2B**). These labeled cells included SG basal layer cells, sebocytes and differentiated cells of the sebaceous duct, but did not include the interfollicular epidermis (IFE), isthmus or other hair follicle compartments. We also did not detect any additional epithelial cell labeling in anagen hair follicles, demonstrating the exquisite specificity for SG labeling in this system (**Figure 2B**).

To track the long-term fate of labeled cells in the SG, we next treated 8 week old mice with doxy-containing chow to suppress any additional new labeling (label-on→off). After 30 weeks of continuous doxy treatment, we observed that ∼86% of SGs were still completely labeled (**Figure 2C-D**). To verify that induction of YFP labeling is indeed suppressed by doxy-chow, we examined adult mice that were continuously treated with doxy since gestation and observed no SG labeling, as expected (**Figure 2C**). These findings suggest that under homeostatic conditions, SGs are largely self-maintained by their own dedicated stem cell pools, but may receive occasional cellular input from the hair follicle.

Following skin injury, stem cells from the IFE and hair follicle migrate into the wound to promote re-epithelialization (Haensel et al., 2020; Huang et al., 2020; Ito et al., 2005; Page et al., 2013; Vagnozzi et al., 2015). To determine whether SG-derived cells exhibit similar behavior, we generated mice with labeled SGs, treated them with doxy to suppress any additional labeling (label-on→off), and subsequently performed excisional wounding. One week after injury, we observed labeled cells that had moved directly out of the SG and into the migratory epithelial front (**Figure 2E**). These SG-derived cells contributed long-term to the regenerated epidermis, since labeled cell clones were still observed up to 8 weeks after wounding (**Figure 2E**). Notably, after skin healing, we observed that only ∼10% of SGs located closest to the wound were labeled, whereas nearly all SGs situated away from the wound were YFP+ (**Figure 2F-G**). This suggests that injury can spur a dramatic reorganization of the SG, where resident SG progenitors depart their niche and are replaced by hair follicle-derived stem cells. The absence of labeling in wound-proximal SGs also provides technical reassurance that *de novo* labeling of SGs is properly suppressed by doxy treatment.

### Sebocyte isolation

Having characterized the cell dynamics of SGs during homeostasis and injury, we next sought to investigate the molecular changes that occur during sebocyte differentiation. While previous scRNA-seq studies on mouse and human skin have included SG subpopulations, these cells are poorly represented due to challenges associated with isolating large, complex, lipid-filled sebocytes. Since PPARγ;YFP mice exhibit specific labeling of SGs, we analyzed skin epithelial cell suspensions by flow cytometry and found that YFP+ cells typically comprise 2-4% of live cells recovered from 8 week old label-on mice (**Figure 3A**). By further fractionating YFP+ cells by size and complexity (measured by forward scatter, FSC, and back scatter, BSC), followed by staining plated cells with the lipophilic dye Nile Red, we observed that the vast majority of Nile Red+ sebocytes were found within the highest ∼10% FSC/BSC subpopulation (**Figure 3A-B**). In contrast, FSC/BSC-low, YFP+ cells were only occasionally stained by Nile Red and likely comprise a mix of smaller SG basal progenitors, early sebocytes and sebaceous duct cells (**Figure 3B**). Overall, this approach enabled us to significantly enrich for SG cells and especially sebocytes, which accounted for < 1% of all cells in our original suspension.

**Figure 3.**
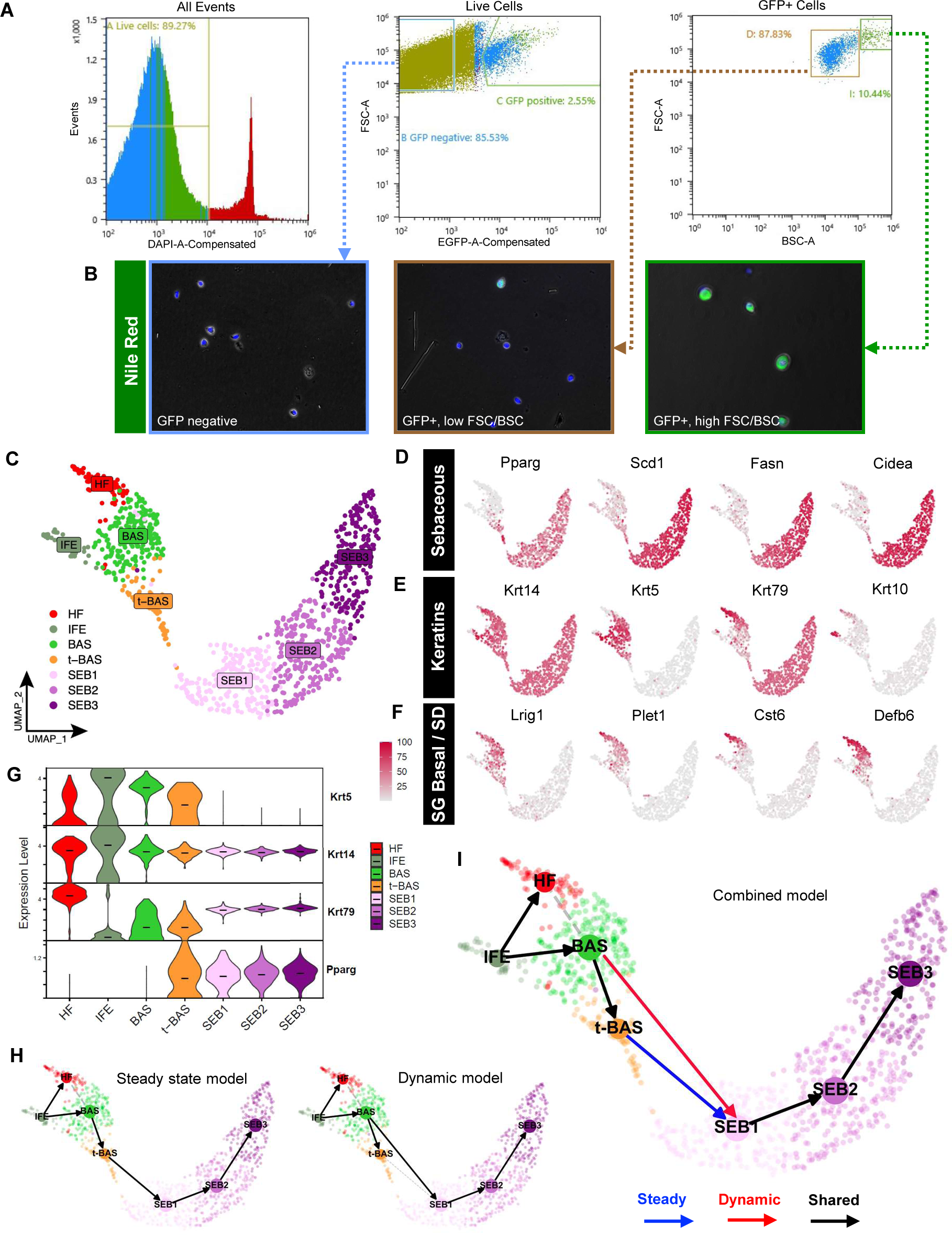
Isolating and profiling SG cells. **A.** Flow cytometry plots of isolated cell suspensions from 8 week old PPARγ;YFP skin. **B.** Nile red staining (green) of sorted keratinocyte subpopulations: bulk GFP negative (left), GFP+ with low FSC/BSC (middle), and GFP+ with high FSC/BSC (right). Note that GFP epifluorescence is not visible and does not interfere with bright Nile red staining, which was superimposed upon bright-field images. **C.** UMAP projection showing 7 cell clusters isolated from YFP-sorted, 8 week old PPARγ;YFP label-on skin. **D.** Feature plots for canonical SG genes. **E.** Feature plots for key keratin genes. **F.** Feature plots for previously identified markers of SG basal progenitors and sebaceous duct (SD). **G.** Violin plots showing relative expression of key marker genes across different cell sub-populations. Note that t-BAS cells uniquely express both *Krt5* and *Pparg*. Horizontal lines indicate median values. **H.** RNA-velocity trajectory analysis performed using either a steady state (left) or dynamic (right) model using scVelo. **I.** Trajectory analysis incorporating results from both steady state and dynamic models, suggesting that BAS cells enter the transitional t-BAS state before differentiating into SEB-1 sebocytes (blue arrow) or can differentiate directly into SEB-1 sebocytes (red arrow). Black arrows, lineage relationships identified by both models.

### Characterizing initial sebocyte differentiation

After devising a strategy to isolate SG cells, we performed targeted scRNA-seq on YFP+ cells sorted from 8 week old skin and visualized these data in 2-dimensional space by uniform manifold approximation and projection (UMAP) using Seurat. We identified 7 cell clusters, including 3 sebocyte clusters (SEB1-3) that exhibit expression of established SG biomarkers (*Pparg, Scd1, Fasn, Cidea*) (**Figure 3C-D**). We also identified 1 cluster representing SG basal cells (BAS) that expresses high level *Krt5, Krt14* and *Lrig1*, which encode markers of SG stem cells (Page et al., 2013) (**Figure 3C, 3E-F**). Flanking the BAS cluster, 1 minor cluster likely comprises differentiated cells of the hair follicle infundibulum and/or sebaceous duct (HF), as assessed by markers *Krt79, Krt17, Krt10, Cst6, Plet1* and *Defb6*, catalogued previously by us and others (**Figure 3C, 3E-F, S1**) (Panteleyev et al., 1997; Raymond et al., 2010; Veniaminova et al., 2013; Zeeuwen et al., 2002). A second minor cluster consists of blended *Krt5*+ basal and *Krt1*+ suprabasal cells of the interfollicular epidermis (IFE), likely originating from SG cells that had departed their niche following mild skin agitation such as scratching (**Figure 3C, S1**).

Notably, a final cell cluster extended out from the BAS cluster towards the SEB sub-populations. Cells in this cluster uniquely express a combination of basal markers (*Krt14, Krt5*) as well as sebocyte differentiation markers (*Pparg*), strongly suggesting that these are the K5+PPARγ+ transitional basal cells (t-BAS) identified above (**Figure 3C, 3G, 1C**). Also consistent with our above findings, we observed that downstream of the t-BAS state, all differentiated SEB clusters express *Krt14* and *Krt79*—but not *Krt5*—reinforcing the notion that SG progenitors undergo a K14:K5 → K14:K79 keratin shift during sebocyte differentiation (**Figure 3G**). Indeed, aside from *Krt14* and *Krt79*, no other keratins were expressed at appreciable levels in the 3 SEB clusters (**Figure S1**), consistent with our previous observation that K79 serves a non-redundant structural role in the SG (Veniaminova et al., 2019).

To infer cell-state transitions between clusters, we next performed RNA-velocity analysis using scVelo and visualized trajectories using partition-based graph abstraction (PAGA). A steady state model of transcriptional dynamics predicted a trajectory whereby BAS cells pass through the t-BAS transitional state to become SEB1 cells (**Figure 3H**, *left*). However, a dynamic model also predicted that a subset of BAS cells can bypass the t-BAS state to directly become SEB1 cells (**Figure 3H**, *right*). Overall, our findings suggest that during homeostasis, resident SG basal progenitors can take either an indirect or direct path to differentiate into SEB1 sebocytes (**Figure 3I**).

### Characterizing sebocyte cell states

Once specified, sebocytes accumulate lipids and undergo a specialized degradative process to release sebum. To better understand the cell-state transitions that occur during sebocyte maturation, we visualized the pseudotemporal dynamics of SG cells using Monocle 2, which ordered the cells in a linear trajectory without significant branching. Although the minor HF and IFE cell states were inter-mixed by this analysis, a single trajectory pointed from BAS to t-BAS, and then sequentially through SEB-1, −2 and −3 terminal states, matching results from RNA-velocity analysis (**Figure 4A-B**).

**Figure 4.**
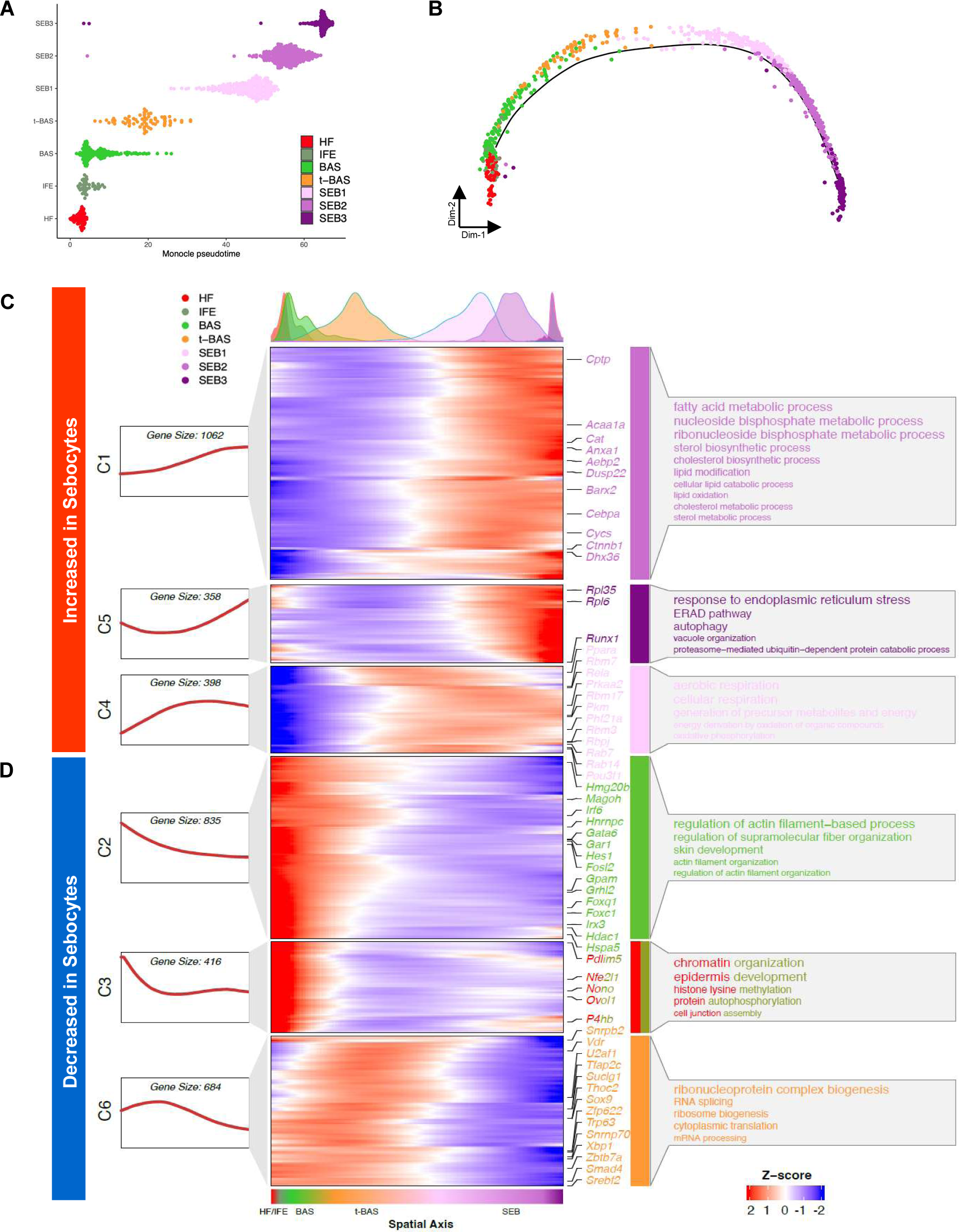
Pseudotemporal dynamics of gene expression during sebocyte differentiation. **A, B.** Pseudotemporal ordering of 7 cell sub-populations isolated from YFP-sorted, PPARγ;YFP label-on skin using Monocle 2. **C, D.** Rolling-wave plot and smoothed expression pattern of pseudotime-dependent genes (n = 3,753) that cluster into six gene modules (C1-C6). Peak positions of the cell populations visualized by kernel density estimation (top), along the pseudospatial axis (bottom). Also shown are the corresponding expression curve (left) and representative enriched GO terms (right) for each gene module. Transcription factors from each module are indicated.

We next identified pseudotime-dependent differentially expressed genes (DEGs) and performed Gene Ontology (GO) analysis to identify cellular processes that become altered during sebocyte maturation. Across the pseudotime trajectory, 3,753 DEGs were identified and grouped by K-medoid clustering into 6 gene modules with distinct expression patterns and biological functions. Notably, 3 modules (C1, C5, C4) of gene expression changes were increased in sebocytes relative to the other cell populations. These modules included genes associated with lipid metabolism, ER stress response, autophagy and aerobic respiration (**Figure 4C**). By contrast, 3 modules (C2, C3, C6) were decreased in sebocytes and were associated with cell functions such as mRNA processing, translation, chromatin organization and cytoskeletal processes (**Figure 4D**). Taken together, these changes confirm that sebocytes comprise a terminally differentiating cell lineage characterized by the shutdown of core cellular processes, the degradation of key structural components, and finally autophagic cell death.

### Spatially mapping sebocyte cell states

Our RNA-velocity and pseudotime analyses both suggest that sebocytes undertake a unidirectional SEB-1 → SEB-2 → SEB-3 trajectory. This path is further supported by a step-wise elevation in expression of canonical SG markers, such as *Scd1, Fasn* and *Mc5r* (**Figure 5A**). To spatially resolve the 3 SEB clusters in the SG, we identified DEGs that define each cell state and found that SEB-1 cells are enriched for *Acp5* and *Mgst2* expression (**Figure 5A, S2**). Although the SEB-2 cluster appears to represent an intermediate state with no unique markers, SEB-3 cells display increased *Awat1* and *Slc6a19* mRNA (**Figure 5A, S2**).

**Figure 5.**
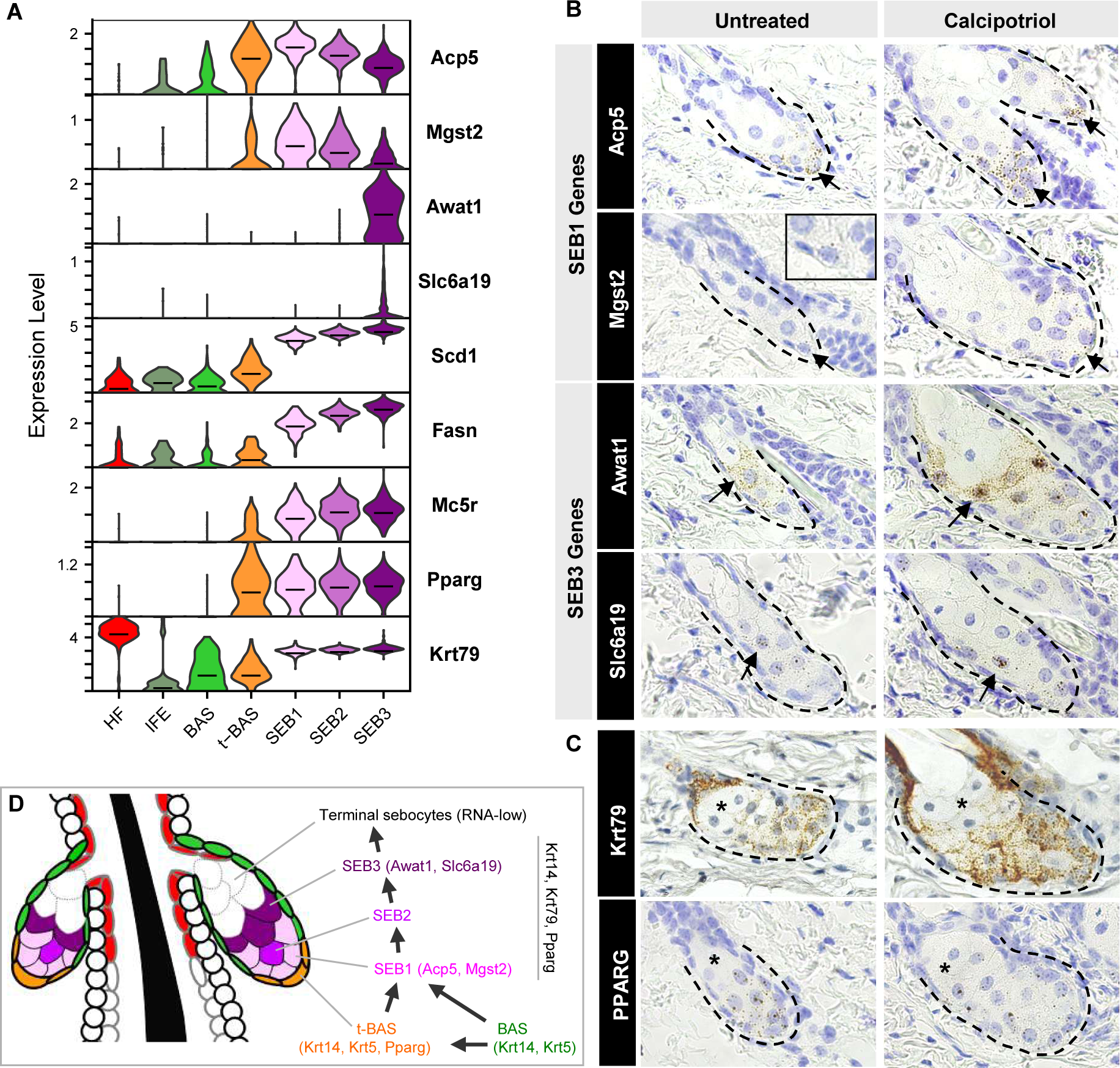
Spatial mapping of different sebocyte cell states. **A.** Violin plots showing relative expression of key marker genes in the SG. Horizontal lines indicate median values. **B.** RNAscope *in situ* staining for genes enriched in SEB-1 (*Acp5, Mgst2*), and genes enriched in SEB-3 (*Awat1, Slc6a19*). Arrow, region where gene is most highly expressed. Inset, magnified view of *Mgst2* staining. **C.** RNAscope staining for highly expressed genes expressed throughout the SG (*Krt79, Pparg*). Asterisk, RNA-low terminal sebocytes. Left column, untreated wild-type skin. Right column, calcipotriol-treated skin. **D.** Schematic summarizing both direct and indirect paths for differentiation of SG basal cells into sebocytes.

By RNAscope *in situ* staining, we next confirmed that expression of *Acp5* and *Mgst2* (SEB-1) is predominantly localized to the lower SG (**Figure 5B**). On the other hand, expression of *Awat1* and *Slc6a19* (SEB-3) is enriched in sebocytes occupying a more central position in the gland (**Figure 5B**). For all 4 genes, we further observed that their expression patterns are recapitulated in skin treated with calcipotriol, a Vitamin D analog that causes SG enlargement, facilitating the visualization of low abundance transcripts (*Mgst2, Slc6a19*) (**Figure 5B**).

Finally, we observed an additional sebocyte population that is rarely stained by any RNAscope probes, including probes targeted against highly expressed pan-SG markers such as *Pparg* and *Krt79* (**Figure 5A, C**). These sebocytes, located at the distal end of the gland, closest to the sebaceous duct, comprise roughly 20-50% of the total SG volume, and likely represent the most terminal cell state downstream of SEB-3. Because these terminal sebocytes are RNA-low, they are likely not represented in our scRNA-seq dataset. Overall, our findings suggest that resident basal progenitors differentiate into SEB1 sebocytes primarily in the lower SG, and that these sebocytes transition unidirectionally along multiple cell states as they move towards the sebaceous duct, as summarized in **Figure 5D**.

### SGs regenerate following genetic ablation

Given that *Pparg* is expressed in both the t-BAS and SEB1-3 cell states (**Figure 5A**), we next tested its requirement for SG homeostasis in adult skin. We therefore generated mice expressing tamoxifen-inducible <u>L</u>rig1-CreERT2 coupled with homozygous conditional alleles for *<u>P</u>parg* (LP mice), which enables targeted deletion of *Pparg* in SG stem cells (He et al., 2003; Powell et al., 2012). When 8 week old LP mice were treated with tamoxifen (TAM)-containing chow for 5 continuous weeks, 99% of SGs were ablated from dorsal skin, confirming the absolute requirement for PPARγ in SG maintenance (**Figure 6A**) (Sardella et al., 2018). Surprisingly, however, when these LP mice were subsequently removed from TAM treatment (“chase”), roughly half of all SGs reappeared within 5 weeks, with full recovery seen after 15 weeks’ chase (**Figure 6A-B**). Regenerated SGs expressed PPARγ, indicating that they were derived from cells that had not undergone Cre-mediated recombination (**Figure S3A**). Since SG regeneration has not been previously documented, these observations propelled our studies in an unexpected direction.

**Figure 6.**
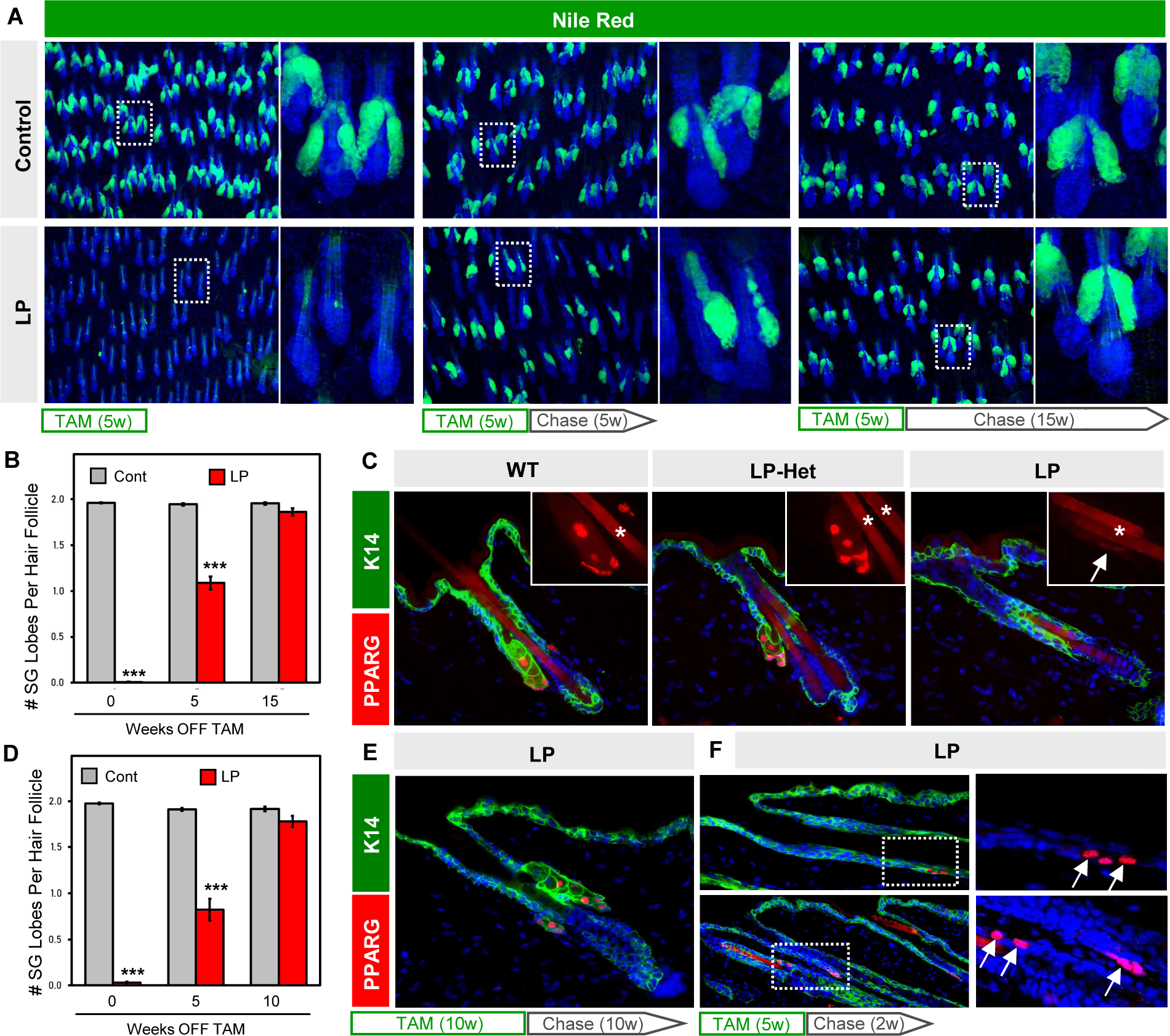
SGs regenerate following genetic ablation. **A.** Nile Red staining (green) of skin whole-mounts from control or LP mice treated with tamoxifen (TAM)-containing chow for 5 continuous weeks, then moved onto normal chow (“chase”) for an additional 0 (left), 5 (middle) or 15 (right) weeks. Right panels are magnified views of the boxed areas. **B.** Quantitation for (A). **C.** Localization of PPARγ (red) in control (left), *Lrig1-CreERT2*;*Pparg-flox*/+ (“LP-Het,” middle) or LP mice (right) following 5 weeks of TAM-chow. Insets, magnified views of PPARγ staining. Arrow, faint PPARγ staining at the hair follicle isthmus in LP skin. Asterisk, hair shaft autofluorescence. **D.** Quantitation of SGs similar to (B), but for mice treated with 10 continuous weeks of TAM-chow, followed by 0-10 weeks’ chase. **E.** Regenerated SGs express PPARγ (red). **F.** Expression of PPARγ (red, arrows) in basal K14+ cells (green) of the upper anagen ORS (top panels) and isthmus (bottom panels), after 5 weeks of TAM-chow and 2 weeks’ chase. Right panels are magnified single channel views of the boxed areas. w, weeks. ***, p < 0.001 by unpaired t-test comparing control and LP skin from the same timepoint.

### Cellular mechanisms for SG regeneration

To better understand how SGs regenerate, we checked whether PPARγ is fully ablated from the hair follicle. In LP mice treated with TAM-chow for 5 weeks (no chase), we observed that PPARγ is almost completely abolished from the isthmus/junctional zone, as expected (**Figure 6C**). However, we also occasionally observed very faint PPARγ staining, at an intensity level far lower than what is seen in skin when only 1 copy of *Pparg* is intentionally deleted (*Pparg*-*flox*/+, or LP-Het) (**Figure 6C**). Thus, trace PPARγ staining in LP follicles is unlikely caused by incomplete recombination within the Lrig1+ domain. Rather, this may reflect cells that had newly entered the isthmus, and had either recently begun expressing PPARγ or had recently deleted PPARγ. Faint PPARγ staining was seen even in LP mice that were treated with TAM-chow for 10 continuous weeks (not shown), and here again PPARγ+ SGs regenerated with similar kinetics after TAM removal (**Figure 6D-E**).

If non-recombined cells enter the isthmus following SG ablation, where are they coming from? To address this, we examined LP mice at earlier timepoints after ceasing TAM treatment. Interestingly, in LP mice treated with TAM-chow for 5 weeks, followed by a shorter 2 weeks’ chase, we observed ectopic PPARγ expression in basal cells within the upper outer root sheath of anagen follicles (**Figure 6F**). This domain has previously been shown to be derived from bulge cells (Hsu et al., 2011), which we confirmed are not targeted by Lrig1-CreERT2 (**Figure S3B**). In addition, we observed high level PPARγ expression reappearing in basal cells at the isthmus, which can also be derived from upper bulge cells over time (**Figure 6F**) (Brownell et al., 2011). Altogether, our findings suggest that non-recombined bulge-derived cells rapidly migrate into the isthmus/junctional zone to regenerate SGs following genetic ablation. In contrast to homeostatic self-renewal, this regenerative process is likely akin to the recruitment of replacement SG progenitors after skin wounding (**Figure 2E-F**).

### Modulation of SG regeneration by hair cycle and FGF signaling

As a final question, we asked what signals instruct progenitor cells to regenerate SGs? For this, we shortened the experimental window and treated 6 week old LP mice with TAM-chow for 2 weeks (no chase), which caused a 97% reduction in PPARγ+/Scd1+ SGs (**Figure 7A, S4A**). At this point, hair follicles have uniformly entered the telogen resting phase at 8 weeks of age. Since subsequent re-entry to anagen growth is asynchronous in adult skin, this provided us the opportunity to assess SG regeneration in anagen and telogen skin from the same animal (**Figure 7B-C**). Indeed, we observed that 5 weeks after TAM withdrawal, anagen skin exhibited a >6-fold increase in SGs compared to adjacent telogen skin from the same animal (**Figure 7C**).

**Figure 7.**
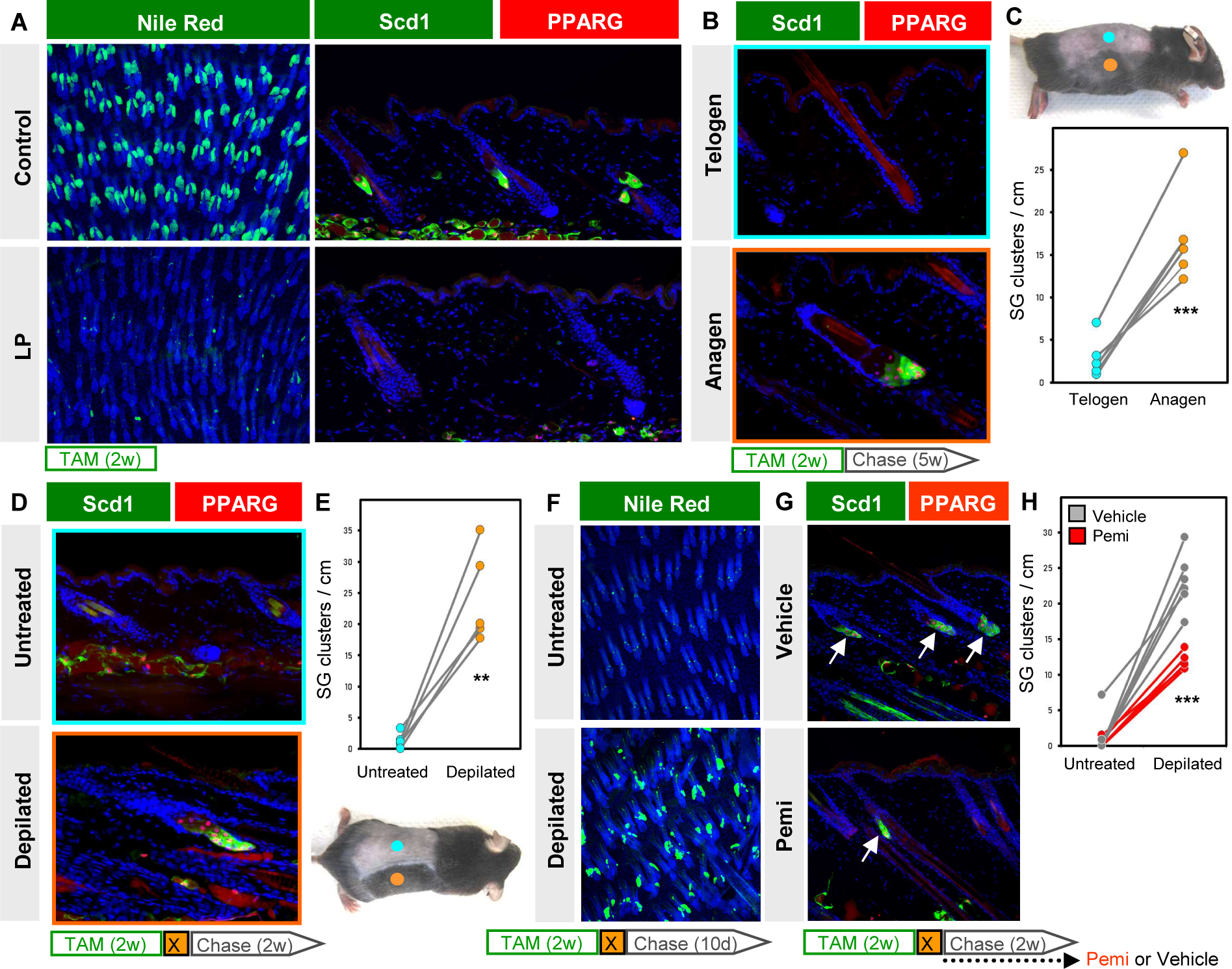
SG regeneration is modulated by hair cycling and FGFR signaling. **A.** Left, Nile Red (green) staining of skin whole-mounts from control (top) or LP (bottom) mice treated with TAM-chow for 2 continuous weeks (no chase). Right, confirmation of SG loss by staining for Scd1 (green) and PPARγ (red). **B.** Scd1/PPARγ staining in telogen (top) or anagen (bottom) skin from the same animal, following 2 weeks of TAM-chow and 5 weeks’ chase. **C.** Top, example of LP mouse treated with TAM-chow for 2 weeks, followed by 5 weeks’ chase. Sites of natural anagen (orange) or telogen (blue) are denoted. Bottom, SG quantitation for (B). Paired samples are connected by lines. **D.** Identification of regenerated SGs by Scd1/PPARγ staining in mice treated with TAM-chow for 2 continuous weeks, then depilated (X) and chased for 2 additional weeks. **E.** Bottom, example of LP mouse used in (D). Sites of depilation (orange) or no treatment (blue) are denoted. Top, quantitation of SG abundance for (D). Paired samples are connected by lines. **F.** Nile Red (green) staining of whole-mounts from untreated (top) or depilated (bottom) LP skin, where mice were treated with TAM-chow for 2 weeks, depilated and chased for 10 days. **G.** Identification of regenerated SGs by Scd1/PPARγ staining (arrows), with similar treatment protocol as in (D), but with additional daily treatment with FGFR inhibitor (pemi) or vehicle during the 2 week chase period. **H.** Quantitation for (G), in LP mice treated with vehicle (grey) or pemi (red). Samples from the same mouse are connected by lines. w, weeks. d, days. **, p < 0.01. ***, p < 0.001. Paired t-test for (C) and (E); unpaired t-test comparing only depilated samples for (H).

To better explore the connection between hair growth and SG regeneration, we next depilated 8 week old LP mice immediately after completing a 2 week course of TAM treatment. Depilation-induced anagen skin similarly exhibited a >10-fold increase in PPARγ+/Scd1+ SGs, compared to non-depilated skin from the same animal (**Figure 7D-E**). These effects were quantitated 2 weeks after depilation/TAM removal, but differences in SG regeneration were apparent even after just 10 days, when most follicles in depilated skin were in early anagen (**Figure 7F**).

Lastly, to identify factors that modulate SG regeneration, we turned back to our scRNA-seq data and found that *Fgfbp3* is among only a handful of genes encoding secreted factors whose expression is enriched in the SEB lineage (**Figure S4B**). Since Fgfbp3 is thought to potentiate FGF signaling (Taetzsch et al., 2018), we treated LP mice with the FGF receptor (FGFR) inhibitor pemigatinib (pemi) (Liu et al., 2020) and examined SG regeneration after depilation. Although FGFR inhibition did not prevent hair follicles from re-entering anagen (**Figure S4C**), significantly fewer SGs regenerated in pemi-treated mice, compared to controls (**Figure 7G-H, S4D**). Altogether, these findings identify a robust and previously unrecognized regenerative process in the skin that is highly influenced by hair growth and FGF signaling, and provides durability in response to injury.

## DISCUSSION

Numerous technical challenges have long hindered the study of SGs. In particular, Cre-mediated approaches for manipulating these appendages typically drive genetic recombination in multiple skin compartments, complicating the interpretation of results. Although mice expressing a sebocyte-specific, *Scd3* promoter-driven Cre have been described, this system likely does not cause recombination in SG basal layer cells, and recombination efficiency in sebocytes remains unclear (Dahlhoff et al., 2016). Another longstanding challenge has been the inability to purify SG cells for molecular profiling. Indeed, we observed that sebocytes constituted <1% of all cells prior to enrichment, consistent with their relative paucity in published scRNA-seq atlases of mouse and human skin (Cheng et al., 2018; Joost et al., 2020; Joost et al., 2016).

By overcoming multiple technical hurdles, our study paints a vibrant portrait of the cellular and molecular architecture of SGs during development, homeostasis, wounding and regeneration. Some themes have emerged: First, SGs are largely self-renewed by resident stem cell pools during homeostasis, although cells originating from outside the gland can also occasionally contribute. Second, when the SG stem cell niche is perturbed, either by wounding or genetic ablation, alternative stem cells rapidly enter the SG domain to repopulate the gland. These findings are consistent with the view that stem cells within discrete hair follicle niches serve largely compartmentalized roles during homeostasis, but become highly plastic following injury (Han et al., 2023; Huang et al., 2020; Schepeler et al., 2014).

A third theme is that while PPARγ is essential for sebocyte differentiation, this transcription factor is initially expressed in SG basal cells. This is seen during development, homeostasis and regeneration. We should emphasize that these t-BAS transitional basal cells—which represent the earliest cells in the SG to express *Pparg*, but also the latest cells to express high level *Krt5* (**Figure 1C, 3G**)—are unlikely to be SG stem cells in adult skin. Similar to transitional basal cells in the interfollicular epidermis that express differentiation markers such as K10, these K5+PPARγ+ cells likely possess limited replication potential and are poised to differentiate (Aragona et al., 2020; Cockburn et al., 2022; Cohen et al., 2022; Wang et al., 2020). While we cannot formally rule out the possibility that t-BAS cells can revert back to PPARγ-negative basal (BAS) cells, which are likely the stem cells that maintain the SG during homeostasis, such a path is not supported by our scRNA-seq trajectory analysis (**Figure 3I**).

If expression of PPARγ indeed predisposes basal cells to differentiate into sebocytes, this raises the question of how the entire SG, including PPARγ-negative basal cells, becomes labeled in adult PPARγ;YFP label-on mice. Unfortunately, examining newborn skin provided little clarity, as early labeling can be seen in both PPARγ+ and PPARγ-cells dispersed around the developing upper follicle prior to formation of the mature SG (**Figure S5A**). Additional studies are needed to clarify how these patterns resolve over time to achieve specific labeling of the entire adult SG. Related to this, we were also unable to acutely switch on labeling of PPARγ+ cells in adult mice that were maintained on doxy-containing chow and subsequently moved onto normal chow (label-off→on) (**Figure S5B**). The reason for this remains unclear; nonetheless, this technical limitation prevented us from tracing the fate of adult PPARγ+ cells.

Previous studies using multi-color lineage tracing have reported that basal cells located along the entire SG periphery can give rise to differentiated sebocytes (Andersen et al., 2019). While our trajectory analyses suggest that BAS progenitors can directly form sebocytes without transitioning through the t-BAS intermediate state, both the direct and indirect paths for sebocyte formation invariably funnel through the SEB-1 state, before moving unidirectionally along progressively more differentiated SEB-2 and SEB-3 lineages. A final, terminal cell state— defined not by scRNA-seq, but instead by low level RNA *in situ* staining—juxtaposes the sebaceous duct (**Figure 5B-C**). Since SEB-1 sebocytes are located at the proximal end of the SG, this implies that new sebocyte formation also primarily occurs within the lower SG. Why our observations differ from those of previous reports remains unclear, but may have to do with the complex geometry of the gland, as well as differences in experimental timing.

Unexpectedly, we observed that SGs regenerate following genetic ablation of PPARγ, and that non-recombined, bulge-derived cells likely give rise to regenerated glands. Although we detected ectopic PPARγ expression in the upper outer root sheath (ORS) of mutant follicles (**Figure 6F**), SGs reappeared at the original sites from where they were lost. These findings demonstrate that bulge-derived cells—which can either move upwards after wounding or downwards during hair growth—have the potential to express PPARγ upon departing their niche. At the same time, the factors that specify the exact site of SG development and regeneration remain elusive. Some of these factors likely involve gradients of Wnt and Hedgehog signaling, as well as AP-1 transcription factor activity, since perturbation of any of these components can drive ectopic SG formation (Allen et al., 2003; Gu and Coulombe, 2008; Petersson et al., 2011; Singh et al., 2018). These gradients may potentially specify both permissive sites for SG formation, as well as non-permissive zones, such as the ORS, which does not form SGs in spite of ectopic PPARγ expression in LP mutants.

Our hair cycle studies also revealed that anagen hair growth, a process associated with increased cell proliferation and movement, greatly accelerates SG regeneration. In contrast, SGs hardly regenerate in telogen skin, indicating that follicles do not automatically regenerate SGs by default. Rather, microenvironmental factors in the skin are likely also critical. At least one of these factors may be Fgfbp3, which binds and liberates FGFs from the extracellular matrix to activate FGF receptors (Taetzsch et al., 2018). Although *Fgfbp3* null mice do not possess obvious skin defects (Tassi et al., 2018), mutant mice lacking Fgfr2 have smaller SGs in tail skin (Grose et al., 2007). Concordantly, acute genetic deletion of *Fgfr2* causes atrophy of eyelid meibomian glands, which are highly related to SGs, and these glands can also partially recover over time (Parfitt et al., 2016; Yang et al., 2021). Other glandular epithelia such as mammary and prostate glands can similarly regenerate after experimental injury in a manner that recapitulates embryonic development (Centonze et al., 2020).

While SG regeneration has not been previously reported, SG loss or hypoplasia has been associated with several skin pathologies including cicatricial alopecia, psoriasis and atopic dermatitis (Karnik et al., 2009; Rittié et al., 2016; Shi et al., 2015; Stenn et al., 1999). Chemotherapy can also induce SG atrophy (Selleri et al., 2006), while lymphocytic attack of SGs has been observed in a mouse model of acute graft versus host disease (Murphy et al., 1991). Isotretinoin, which is used to treat severe acne, reduces SG size by up to 90% (Goldstein et al., 1982; Landthaler et al., 1980). Even in normal skin, SG size and activity increase and diminish at different stages throughout life (Plewig and Kligman, 1978; Zouboulis and Boschnakow, 2001). Whether SGs undergo regeneration in these varied contexts remains unclear but is conceivable in light of our findings. In summary, our work identifies distinct cellular mechanisms for SG maintenance and regeneration, which may ultimately enable these appendages to be preserved following challenges to the skin.

## ACKNOWLEDGEMENTS

We are grateful to the Dlugosz lab (University of Michigan) for helpful discussions and sharing reagents; Dr. Y. Eugene Chen (University of Michigan) for sharing mice; Ann Marie Deslauriers-Cox for flox cytometry; Dr. Allison C. Billi (University of Michigan) for advice on single cell preparation; and Olivia Koues, Tricia Tamsen and the Advanced Genomics Core (University of Michigan) for single cell sequencing. S.Y.W. acknowledges the support of the Leo Foundation (LF18017) and the NIH (R01AR065409 and R01AR080654). S.X.A. acknowledges the support of the National Science Foundation (CBET2134916). S.Y.W. and S.X.A. were jointly funded by the American Cancer Society (TLC-21-161-01-TLC). The authors also acknowledge support from the UM Skin Biology and Disease Resource-based Center (P30AR075043) and NCI Cancer Center Support Grant (P30CA046592).

## AUTHOR CONTRIBUTIONS

Conceptualization and methodology, N.A.V., Y.J, A.A.D., S.X.A., S.Y.W.

Investigation, N.A.V., A.H., T.J.H., S.Y.T., M.G., S.N., S.Y.W.

Formal Analysis, N.A.V., Y.J., S.X.A., S.Y.W.

Writing – Original Draft, Review & Editing, N.A.V., Y.J., S.X.A., S.Y.W.

Funding Acquisition and Supervision, S.X.A., S.Y.W.

## SUPPLEMENTAL FIGURE LEGENDS

**Figure S1.** Feature plots for 28 keratin genes. Only keratin genes that met filtering criteria are depicted.

**Figure S2.** Top 10 genes enriched in each cell cluster, calculated by COSG (A) or Seurat-Wilcoxon (B), depicted by heat maps (left) or dot plots (right). Red denotes high expression, blue denotes low expression.

**Figure S3. A.** Regenerated SG (right hair follicle) has PPARγ+ cells (red), following 5 weeks of TAM chow treatment and 5 weeks’ chase. **B.** Lrig1-CreERT does not induce recombination, as assessed by YFP reporter expression (green), in the hair follicle bulge (arrow), following 5 weeks of TAM chow treatment.

**Figure S4. A.** Quantitation of SG abundance in LP mice treated for 2 weeks with TAM chow beginning at 6 weeks of age. SGs were counted in frozen skin sections stained for PPARγ and Scd1. Control animals were Cre-negative littermates treated with TAM chow. **B.** Feature plot for *Fgfbp3*. **C.** Skin from LP mice that were depilated (boxed areas) and treated systemically with either vehicle or pemi during the 2 week chase period entered anagen and regrew hair normally. **D.** Quantitation of SG abundance in LP mice treated with 2 weeks of TAM chow beginning at 6 weeks of age, then depilated, and treated with either vehicle (grey) or pemi (red) during the 2 week chase period. Paired samples collected from the same mouse are connected by lines. Quantitation was performed on skin from the same animals as shown in Figure 7H, using an alternative quantitation method where SGs were counted from H&E-stained sections. **, p < 0.01 by unpaired t-test comparing depilated pemi-treated versus depilated vehicle-treated samples.

**Figure S5. A.** Skin from PPARγ;YFP label-on mice, harvested at 1 week (left) or 2 weeks (right) of age. Note the heterogeneous recombination, as assessed by YFP reporter expression (green), in PPARγ+ cells (red) as well as PPARγ-negative cells. Lower panels show identical views as above, with DAPI omitted. **B.** Skin from PPARγ;YFP mice that were treated with doxy since gestation until 8 weeks of age (label-off), then followed for an additional 6 weeks (left) or 12 weeks (right) after doxy removal (label-off→on). Note that this approach does not cause YFP reporter expression in this system.

**Figure S6. A.** Quality control metrics of 4 replicate single cell libraries (LT1-4) before threshold filtering. **B.** Metrics of single cell libraries after threshold filtering. Cells with <200 detected genes as well as those with >15% mitochondrial content were removed. Cells having >7,000 detected genes were removed to exclude potential doublets. Cells with <900 and >80000 UMIs per cell were removed. Genes detected in <3 cells were removed. **C.** Each library was projected by UMAP and found not to exhibit significant batch effects. **D.** PCA analysis combining all 4 replicates. **E.** PC1 score distribution by VlnPlot across 4 replicates.

**Figure S7. A.** Cell cycle annotation performed using the Cell Cycle Scoring function in Seurat. **B.** Cycling cell scoring. A set of S- and G2M-phase genes was combined and used for cycling cell scoring, which was calculated by the AddModuleScore function with default parameters in Seurat. **C.** Cycling cell scoring across different cell states. Dashed line indicates the mean score across all cells. **D.** Cell type signature scoring calculated by UCell to identify cell populations (see Methods for details). Key genes for each cell type used in the signature are shown.

## MATERIALS AND METHODS

### Lead Contact

Further information and requests for resources and reagents should be directed to and will be fulfilled by the Lead Contact, Sunny Wong.

### Materials Availability

All reagents generated in this study are available from the lead contact.

### Drug treatment

For labeling studies, PPARγ;YFP mice were fed doxycycline-containing chow (1 g/kg, BioServ Inc, F3949) ad libitum to suppress tTA activity (label-off). For SG ablation and regeneration studies, LP mice and Cre-negative littermate controls were fed irradiated TAM-containing chow (400 mg/kg, Envigo TD.130860). Pemigatinib (INCB054828, SelleckChem) was dissolved in DMSO to a stock concentration of 4 mg/mL, then subsequently diluted in PEG 400/5% dextrose in water (75:25 v/v). Mice were treated daily at a dose of 1 mg/kg body weight by oral gavage for 14 consecutive days after depilation during the chase period. Calcipotriol (C4369, Sigma) was dissolved in 100% ethanol and 5.3 nmols were applied topically onto shaved skin for 9 consecutive days at a volume of 200 μL, then harvested 1 day after the final treatment.

### Whole mount analysis

Whole mounts of telogen dorsal skin were performed as previously described (Veniaminova et al., 2019). Briefly, skin was shaved, excised, stretched on a paper towel, covered with Elmer’s No-Wrinkle rubber cement and overlayed with cellophane tape. Following incubation for 6 hours in 5 mM EDTA/PBS at 37 degrees Celsius, the epidermis was separated from the dermis and fixed in 3.7% formalin for 30 minutes at room temperature. Finally, the samples were incubated with Nile Red (4 μg/ml) and DAPI (1 μg/ml) for 30 min in PBS with gentle agitation at room temperature, then mounted with Vectashield on a microscope slide and imaged.

### Flow cytometry

Label-on PPARγ;YFP mice were euthanized at 8 weeks of age, and dorsal skin was shaved and removed. The epidermis was separated from the dermis and cell suspensions were obtained by overnight trypsinization (0.25% trypsin, Invitrogen) at 4 degrees Celsius, as previously described (Veniaminova et al., 2013). Single cells were resuspended in 2% BSA/HBSS, stained with DAPI to exclude dead cells, and sorted using a SH800 cell sorter (Sony). For scRNA-seq, 60,000 YFP+ cells from an 8 week-old PPARγ;YFP label-on male mouse were sorted into 2% BSA/HBSS buffer, at a ratio of 3:1 FSC/BSC-high:FSC/BSC-low, where “high” cells represented the largest ∼10% of cells by FSC/BSC, and “low” cells comprised the remaining 90% by FSC/BSC (Figure 3A).

### Single cell library preparation

Single cell suspensions were subjected to counting on the LUNA Fx7 Automated Cell Counter (Logos Biosystems) and diluted to a concentration of 300 cells/μL. Single nuclei 3’ Gene Expression LT libraries were generated using the 10x Genomics Chromium instrument following the manufacturer’s protocol (Chromium Next GEM Single Cell 3’ LT Kit v3.1). In brief, suspensions were loaded onto the 10x chip along with reverse transcription (RT) master mix and appropriate gel beads. Following generation of single-cell gel bead-in-emulsions (GEMs), reverse transcription was performed, and the resulting Post GEM-RT product was cleaned up and the cDNA is amplified. cDNA was subjected to enzymatic fragmentation and size selection to optimize the cDNA size prior to final library construction following the manufacturer’s protocol (10x Genomics). Final library quality was assessed using the LabChip GX (PerkinElmer). Libraries were then subjected to paired-end sequencing according to the manufacturer’s protocol (Illumina NovaSeq 6000). Four LT reactions were run in parallel (LT1-4) from the same animal.

### Single cell data processing and analysis

Bcl2fastq2 Conversion Software (Illumina) was used to generate de-multiplexed fastq files, and the CellRanger Pipeline (10x Genomics) was used to align reads and generate count matrices against the mouse genome GRCm38/mm10. For downstream analysis, the Seurat (v4.3.0) R package (Hao et al., 2021) was used to combine the 4 cell libraries and a merged Seurat object was generated. Genes detected in <3 cells were removed. Low-quality cells were further filtered on the basis of total UMI counts per cell (>900 and <80,000), number of detected genes (>200 and <7,000) and mitochondrial genes fraction (<15%). Applying these filters resulted in a final dataset of 1,066 single cell transcriptomes (**Figure S6A-B**).

To account for batch effects, the merged Seurat object was normalized using the NormalizeData() function with a scale factor of 10,000, and variable features were identified using FindVariableFeatures() with 2,000 genes. Principal component analysis (PCA) was used and the first 30 principal components (PCs) were further summarized using UMAP dimensionality reduction. We chose to use 30 PCs based on results from analyses using Elbow plots. Clustering was conducted using the FindNeighbors() and FindClusters() functions using 30 PCA components and a resolution parameter set to 0.7. A library-split UMAP plot was generated by DimPlot() function to evaluate inter-sample differences. For batch effect detection across different libraries, the distribution of the first principal component (PC1) obtained after PCA analysis was visualized by VlnPlot(). As no obvious batch effect was observed between samples (**Figure S6C-E**), we utilized the processed merged Seurat object for subsequent analysis.

For potential doublet detection, we identified doublets with DoubletFinder (v2.0) (McGinnis et al., 2019). The doublets were predicted using the cleaned pre-processed merged Seurat data. We did not filter doublets because no discrete doublet-enriched cluster was identified, and only few doublets were observed in the dataset.

Cluster markers were interpreted and assigned using established cell type annotations: *Krt5*/*Krt1*(+), *Lrig1*(-) and *Pparg*(-) for blended interfollicular epidermis (IFE); *Defb6*(+), *Cst6*(+), *Krt17*(+) and *Krt79*(+) for hair follicle-related cells (HF); *Krt5*(+), *Krt14*(+), *Lrig1*(+) and *Pparg*(-) for SG basal cells (BAS); *Krt5*(+) and *Pparg*(+) for transitional basal cells (t-BAS); and *Cidea*(+), *Scd1*(+) and *Fasn*(+) for differentiated sebocytes (SEB1/2/3).

To assess the effects of cell cycle heterogeneity on cell clustering, cell cycle phase scores were estimated using Seurat’s CellCycleScoring function with mouse homologs of the cell cycle gene sets provided by Seurat. No obvious clustering differences were found between G2M and S phases within differentiating cells (**Figure S7A-B**). The signals separating non-cycling cells and cycling cells were also checked by combined G2M and S phase gene scoring (cycling cell scoring) and showed high correlation between cycling cell score and corresponding cell states (**Figure S7C**).

To identify DEGs in each cell cluster, we used the Seurat FindAllMarkers function and the COSG (v0.9.0) R package (Dai et al., 2022) (**Figure S2**). The COSG-identified top genes were used to establish the cell identity of each cluster, along with markers described in the literature for assigned cell states. Gene signature scores were calculated on the basis of the scRNA-seq data. For each gene signature, individual cells were scored using UCell (v2.2.0) R package (Andreatta and Carmona, 2021) and projected onto UMAP plots (**Figure S7D**).

scVelo (v.0.2.5) (Bergen et al., 2020) and Monocle 2 (Qiu et al., 2017a; Qiu et al., 2017b) were used for trajectory analysis. For scVelo, reads that passed quality control after clustering were used as input for the velocyto command line. The mouse expressed repeat annotation file was retrieved from UCSC genome browser. The genome annotation file was provided by CellRanger. The output loom file was used as input to estimate velocity. Velocity embedding was estimated using either the steady-state or likelihood-based dynamical model. PAGA was performed using the sc.tl.paga function in scVelo. For Monocle 2, we built a new CellDataSet object from the cluster-annotated Seurat object using the newCellDataSet function. We used the differentialGeneTest function to derive DEGs from each cluster, and genes with q[<[1[×[10^−4^ were used to order cells in pseudotime. Dimension reduction was performed using the DDRTree algorithm and cells were ordered along the trajectory. Gene Ontology enrichment analysis was performed using clusterProfiler (Yu et al., 2012). bitr() was first employed to map gene symbols to Entrez IDs using org.Mm.eg.db (v3.16.0) (Carlson, 2015) as the reference database, and then the enrichGO function was used with “ont[=[“BP”, pAdjustMethod[=[“BH”, pvalueCutoff [=[0.01, and qvalueCutoff[=[0.05”.

### Immunohistochemistry and RNAscope

Frozen sections were probed with antibodies against the following antigens: GFP (1:1000), Ki67 (1:300), K14 (1:1000), K5 (1:1000), K79 (1:400), PPARγ (1:300) and Scd1 (1:300). In some cases, fluorescent images were processed using the Auto-Blend feature of Adobe Photoshop CS6 to automatically maximize image sharpness across multiple focal planes. RNAscope 2.5 Brown kit (ACD Bio) was used for RNA *in situ* staining according to manufacture’s protocol. After deparaffinization, 5 μm sections were boiled for 15 minutes in RNAscope retrieval buffer, treated with protease for 30 minutes and incubated with target probes for 2 hours at 40 degrees Celsius. Probe detection was performed according to manufacturer’s instructions.

### SG Quantitation

All analyses utilized a minimum of 3 mice per gender, genotype and timepoint. Experiments utilized matched mutant and control litter-mate animals, whenever possible. To quantitate SGs in whole mounts, 2 representative fields at 5x magnification were photographed for DAPI and Nile Red staining, and subsequently all images were divided into thirds by drawing guide lines. SG presence or absence was scored for every third hair follicle that intersected these guide lines, yielding 18-25 randomly selected follicles per field. To quantitate SGs from sections, frozen skin sections (8 μm) were stained with antibodies against PPARγ and Scd1. The number of PPARγ/Scd1 double-positive SG clusters was then counted and normalized to the length of the skin section.

### Statistics

SG quantitation data are depicted as means from independent biological replicates. Unpaired t tests were performed in most cases to determine statistical significance. For matched samples harvested from the same animal, paired t tests were used for comparisons between groups. Error bars are depicted as SEM.

| REAGENTS or RESOURCES | SOURCE | IDENTIFIER |
| --- | --- | --- |
| **Antibodies** |  |  |
| Chicken anti-K14 | Biolegend | Cat # 906004 |
| Chicken anti-K5 | Biolegend | Cat # 905903 |
| Rat anti-Ki67 | eBioscience | Cat # 14-5698-80 |
| Goat anti-K79 | Santa Cruz | Cat # sc-243156 |
| Rabbit anti-PPARγ | Cell Signaling | Cat # 2443S |
| Chicken anti-GFP | Abcam | Cat # ab13970 |
| Goat anti-Scd1 | Santa Cruz | Cat # sc-14719 |
| **Biological samples** |  |  |
| Mouse tissue samples, obtained in accordance with guidelines established by the University of Michigan Unit for Laboratory Animal Medicine | This manuscript | Study protocol # PRO00010041 |
| **Chemicals, peptides, and recombinant proteins** |  |  |
| Doxycycline chow (1 g/kg) | BioServ Inc | Cat # F3949 |
| Tamoxifen chow (400 mg/kg, irradiated) | Envigo | Cat # TD.130860 |
| 0.25% Trypsin (no EDTA) | Invitrogen | Cat # 15050065 |
| Albumin, Bovine Fraction V (BSA) | Research Prod. International | Cat # A30075 |
| Hank's balanced salt solution (HBSS) | Gibco | Cat # 14025092 |
| Nair hair removal lotion | Nair | Cat # B001E6OAM8 |
| Hematoxylin | Sigma | Cat # HHS16 |
| DAPI | Sigma | Cat # 32670 |
| Nile Red | Sigma | Cat # N3013 |
| Pemigatinib (INCB054828) | SelleckChem | Cat # S0088 |
| Calcipotriol | Sigma | Cat # C4369 |
| **Critical commercial assays** |  |  |
| RNAscope 2.5 HD Reagent Kit-BROWN | ACD (RNAscope) | Cat # 322310 |
| RNAscope 2.5 Pretreat Reagents-H202 and Protease Plus | ACD (RNAscope) | Cat # 322330 |
| RNAscope Target Retrieval | ACD (RNAscope) | Cat # 322000 |
| RNAscope Wash Buffer | ACD (RNAscope) | Cat # 310091 |
| **Experimental models: Organisms/strains** |  |  |
| Mouse: Lrig1<sup>tm1.1(cre/ERT2)Rjc</sup> (Lrig1-Cre<sup>ERT2</sup>) | The Jackson Laboratory | Cat # 018418 |
| Mouse: B6.129-Pparg<sup>tm2Rev/J</sup> (Pparg-flox) | The Jackson Laboratory<br>(by way of Dr. Y. Eugene Chen) | Cat # 004584 |
| Mouse: B6;129-Pparg<sup>tm1.1(tTA)/Jmrg/J</sup> (AdipoTrak) | The Jackson Laboratory | Cat # 024755 |
| Mouse: B6.Cg-Tg(tetO-cre)1Jaw/J (TRE-Cre) | The Jackson Laboratory<br>(by way of Dr. A. Dlugosz) | Cat # 006234 |
| Mouse: Gt(ROSA)26Sor<sup>tm1(EYFP)Cos</sup> (YFP reporter) | The Jackson Laboratory | Cat # 006148 |
| **Oligonucleotides** |  |  |
| *In situ* probe: mouse Acp5 | ACD (RNAscope) | Cat # 465001 |
| *In situ* probe: mouse Mgst2 | ACD (RNAscope) | Cat # 819931 |
| *In situ* probe: mouse Awat1 | ACD (RNAscope) | Cat # 1172821-C1 |
| *In situ* probe: mouse Slc6a19 | ACD (RNAscope) | Cat # 897821 |
| *In situ* probe: mouse Pparg | ACD (RNAscope) | Cat # 418821 |
| *In situ* probe: mouse Krt79 | ACD (RNAscope) | Cat # 436201 |
| **Software and algorithms** |  |  |
| Cell Ranger v6.1.2 | 10X Genomics | https://support.10xgenomics.com/single-cell-gene-expression/software/pipelines/latest/installation |
| DoubletFinder v2.0 | McGinnis et al. | https://github.com/chris-mcginnis-ucsf/DoubletFinder |
| Seurat v4.3.0 | Hao et al. | https://github.com/satijalab/seurat |
| COSG v0.9.0 | Dai et al. | https://github.com/genecell/COSGR |
| scVelo v0.2.5 | Bergen et al. | https://github.com/theislab/scvelo |
| Monocle 2 | Qiu et al. | https://github.com/cole-trapnell-lab/monocle2-rge-paper |
| clusterProfiler v4.6.0 | Yu et al. | https://github.com/YuLab-SMU/clusterProfiler |
| UCell 2.2.0 | Andreatta et al. | https://github.com/carmonalab/UCell |
| R | R Core | https://www.r-project.org/ |
| Python | Python Software Foundation | https://www.python.org/ |
| org.Mm.eg.db | Carlson | http://bioconductor.org/packages/org.Mm.eg.db/ |

## REFERENCES

Ahmadian, M., Suh, J.M., Hah, N., Liddle, C., Atkins, A.R., Downes, M., and Evans, R.M. (2013). PPARgamma signaling and metabolism: the good, the bad and the future. Nat Med 19, 557–566.

Allen, M., Grachtchouk, M., Sheng, H., Grachtchouk, V., Wang, A., Wei, L., Liu, J., Ramirez, A., Metzger, D., Chambon, P., et al. (2003). Hedgehog signaling regulates sebaceous gland development. Am J Pathol 163, 2173–2178.

Andersen, M.S., Hannezo, E., Ulyanchenko, S., Estrach, S., Antoku, Y., Pisano, S., Boonekamp, K.E., Sendrup, S., Maimets, M., Pederson, M.T., et al. (2019). Tracing the cellular dynamics of sebaceous gland development in normal and perturbed states. Nat Cell Biol 21, 924–932.

Andreatta, M., and Carmona, S.J. (2021). UCell: Robust and scalable single-cell gene signature scoring. Comput Struct Biotechnol J 19, 3796–3798.

Aragona, M., Sifrim, A., Malfait, M., Song, Y., Herck, J.V., Dekoninck, S., Gargouri, S., Lapouge, G., Swedlund, B., Dubois, C., et al. (2020). Mechanisms of stretch-mediated skin expansion at single-cell resolution. Nature 584, 268–273.

Bergen, V., Lange, M., Peidli, S., Wolf, F.A., and Theis, F.J. (2020). Generalizing RNA velocity to transient cell states through dynamical modeling. Nat Biotechnol 38, 1408–1414.

Brownell, I., Guevara, E., Bai, C.B., Loomis, C.A., and Joyner, A.L. (2011). Nerve-derived sonic hedgehog defines a niche for hair follicle stem cells capable of becoming epidermal stem cells. Cell Stem Cell 8, 552–565.

Carlson, M. (2015). Org. Mm. Eg. Db: Genome Wide Annotation for Mouse. Bioconductor, http://bioconductor.org/packages/org.Mm.eg.db/.

Centonze, A., Lin, S., Tika, E., Sifrim, A., Fioramonti, M., Malfait, M., Song, Y., Wuidart, A., Herck, J.V., Dannau, A., et al. (2020). Heterotypic cell-cell communication regulates glandular stem cell multipotency. Nature 584, 608–613.

Cheng, J.B., Sedgewick, A.J., Finnegan, A.I., Harirchian, P., Lee, J., Kwon, S., Fassett, M.S., Golovato, J., Gray, M., Ghadially, R., et al. (2018). Transcriptional programming of normal and inflamed human epidermis at single-cell resolution. Cell Rep 25, 871–883.

Choa, R., Tohyama, J., Wada, S., Meng, H., Hu, J., Okumura, M., May, R.M., Robertson, T.F., Pai, R.L., Nace, A., et al. (2021). Thymic stromal lymphopoietin induces adipose loss through sebum hypersecretion. Science 373, eabd2893.

Cockburn, K., Annusver, K., Gonzalez, D.G., Ganesan, S., May, D.P., Mesa, K.R., Kawaguchi, K., Kasper, M., and Greco, V. (2022). Gradual differentiation uncoupled from cell cycle exit generates heterogeneity in the epidermal stem cell layer. Nat Cell Biol, doi: 10.1038/s41556-41022-01021-41558.

Cohen, E., Johnson, C., Redmond, C.J., Nair, R.R., and Coulombe, P.A. (2022). Revisiting the significance of keratin expression in complex epithelia. J Cell Sci 135, jcs260594.

Cottle, D.L., Kretzschmar, K., Schweiger, P.J., Quist, S.R., Gollnick, H.P., Natsuga, K., Aoyagi, S., and Watt, F.M. (2013). c-MYC-induced sebaceous gland differentiation is controlled by an androgen receptor/p53 axis. Cell Rep 3, 427–441.

Dahlhoff, M., Camera, E., Schäfer, M., Emrich, D., Riethmacher, D., Foster, A., Paus, R., and Schneider, M.R. (2016). Sebaceous lipids are essential for water repulsion, protection against UVB-induced apoptosis, and ocular integrity in mice. Development 143, 1823–1831.

Dai, M., Pei, W., and Wang, X.J. (2022). Accurate and fast cell marker gene identification with COSG. Brief Bioinform 23, bbab579.

Epstein, E.H., and Epstein, W.L. (1966). New cell formation in human sebaceous glands. J Invest Dermatol 46, 453–458.

Feldman, A., Mukha, D., Maor, I.I., Sedov, E., Koren, E., Yosefzon, Y., Shlomi, T., and Fuchs, Y. (2019). Blimp1+ cells generate functional mouse sebaceous gland organoids in vitro. Nat Commun 10, 2348.

Fischer, H., Fumicz, J., Rossiter, H., Napirei, M., Buchberger, M., Tschachler, E., and Eckhart, L. (2017). Holocrine secretion of sebum is a unique DNase2-dependent mode of programmed cell death. J Invest Dermatol 137, 587–594.

Frances, D., and Niemann, C. (2012). Stem cell dynamics in sebaceous gland morphogenesis in mouse skin. Dev Biol 363, 138–146.

Füllgrabe, A., Joost, S., Are, A., Jacob, T., Sivan, U., Haegebarth, A., Linnarsson, S., Simons, B.D., Clevers, H., Toftgård, R., et al. (2015). Dynamics of Lgr6+ Progenitor Cells in the Hair Follicle, Sebaceous Gland, and Interfollicular Epidermis. Stem Cell Reports 5, 843–855.

Ghazizadeh, S., and Taichman, L.B. (2001). Multiple classes of stem cells in cutaneous epithelium: a lineage analysis of adult mouse skin. EMBO J 20, 1215–1222.

Goldstein, J.A., Comite, H., Mescon, H., and Pochi, P.E. (1982). Isotretinoin in the treatment of acne: histologic changes, sebum production, and clinical observations. Arch Dermatol 118, 555–558.

Grose, R., Fantl, V., Werner, S., Chioni, A.M., Jarosz, M., Rudling, R., Cross, B., Hart, I.R., and Dickson, C. (2007). The role of fibroblast growth factor receptor 2b in skin homeostasis and cancer development. EMBO J 26, 1268–1278.

Gu, L.H., and Coulombe, P.A. (2008). Hedgehog signaling, Keratin 6 induction, and sebaceous gland morphogenesis. Am J Pathol 173, 752-761.

Haensel, D., Jin, S., Sun, P., Cinco, R., Dragan, M., Nguyen, Q., Cang, Z., Gong, Y., Vu, R., MacLean, A.L., et al. (2020). Defining epidermal basal cell states during skin homeostasis and wound healing using single-cell transcriptomics. Cell Rep 30, 3932–3947.

Han, J., Lin, K., Choo, H., Chen, Y., Zhang, X., Xu, R.H., Wang, X., and Wu, Y. (2023). Distinct bulge stem cell populations maintain the pilosebaceous unit in a β-catenin-dependent manner. iScience 26, 105805.

Hao, Y., Hao, S., Andersen-Nissen, E., 3rd, W.M.M., Zheng, S., Butler, A., Lee, M.J., Wilk, A.J., Darby, C., Zager, M., et al. (2021). Integrated analysis of multimodal single-cell data. Cell 184, 3573-3587.

He, W., Barak, Y., Hevener, A., Olson, P., Liao, D., Le, J., Nelson, M., Ong, E., Olefsky, J.M., and Evans, R.M. (2003). Adipose-specific peroxisome proliferator-activated receptor gamma knockout causes insulin resistance in fat and liver but not in muscle. Proc Natl Acad Sci USA 100, 15712–15717.

Hinde, E., Haslam, I.S., Schneider, M.R., Langan, E.A., Kloepper, J.E., Schramm, C., Zouboulis, C.C., and Paus, R. (2013). A practical guide for the study of human and murine sebaceous glands in situ. Exp Dermatol 22, 631–637.

Hsu, Y.C., Pasolli, H.A., and Fuchs, E. (2011). Dynamics between stem cells, niche, and progeny in the hair follicle. Cell 144, 92–105.

Huang, S., Kuri, P., Aubert, Y., Brewster, M., Li, N., Farrelly, O., Rice, G., Bae, H., Prouty, S., Dentchev, T., et al. (2020). Lgr6 marks epidermal stem cells with a nerve-dependent role in wound re-epithelialization. Cell Stem Cell 28, 1582–1596.e1586.

Ito, M., Liu, Y., Yang, Z., Nguyen, J., Liang, F., Morris, R.J., and Cotsarelis, G. (2005). Stem cells in the hair follicle bulge contribute to wound repair but not to homeostasis of the epidermis. Nat Med 11, 1351–1354.

Joost, S., Annusver, K., Jacob, T., Sun, X., Dalessandri, T., Sivan, U., Sequeira, I., Sandberg, R., and Kasper, M. (2020). The molecular anatomy of mouse skin during hair growth and rest. Cell Stem Cell 26, 441–457.

Joost, S., Zeisel, A., Jacob, T., Sun, X., Manno, G.L., Lonnerberg, P., Linnarsson, S., and Kasper, M. (2016). Single-cell transcriptomics reveals that differentiation and spatial signatures shape epidermal and hair follicle heterogeneity. Cell Syst 3, 221–237.

Jung, Y., Tam, J., Jalian, H.R., Anderson, R.R., and Evans, C.L. (2015). Longitudinal, 3D in vivo imaging of sebaceous glands by coherent anti-stokes Raman scattering microscopy: normal function and response to cryotherapy. J Invest Dermatol 135, 39-44.

Karnik, P., Tekeste, Z., McCormick, T.S., Gilliam, A.C., Price, V.H., Cooper, K.D., and Mirmirani, P. (2009). Hair follicle stem cell-specific PPARgamma deletion causes scarring alopecia. J Invest Dermatol 129, 1243–1257.

Kobayashi, T., Voisin, B., Kim, D.Y., Kennedy, E.A., Jo, J.H., Shih, H.Y., Truong, A., Doebel, T., Sakamoto, K., Cui, C.Y., et al. (2019). Homeostatic Control of Sebaceous Glands by Innate Lymphoid Cells Regulates Commensal Bacteria Equilibrium. Cell 176, 982–997.

Kretzschmar, K., Cottle, D.L., Donati, G., Chiang, M.F., Quist, S.R., Gollnick, H.P., Natsuga, K., Lin, K.I., and Watt, F.M. (2014). BLIMP1 is required for postnatal epidermal homeostasis but does not define a sebaceous gland progenitor under steady-state conditions. Stem Cell Reports 3, 620–633.

Landthaler, M., Kummermehr, J., Wagner, A., and Plewig, G. (1980). Inhibitory effects of 13-cis-retinoic acid on human sebaceous glands. Arch Dermatol Res 269, 297–309.

Lee, S.J., Seok, J., Jeong, S.Y., Park, K.Y., Li, K., and Seo, S.J. (2016). Facial Pores: Definition, Causes, and Treatment Options. Dermatol Surg 42, 277–285.

Liu, P.C.C., Koblish, H., Wu, L., Bowman, K., Diamond, S., DiMatteo, D., Zhang, Y., Hansbury, M., Rupar, M., Wen, X., et al. (2020). INCB054828 (pemigatinib), a potent and selective inhibitor of fibroblast growth factor receptors 1, 2, and 3, displays activity against genetically defined tumor models. PLoS One 15, e0231877.

Lovászi, M., Szegedi, A., Zouboulis, C.C., and Törőcsik, D. (2017). Sebaceous-immunobiology is orchestrated by sebum lipids. Dermatoendocrinol 9, e1375636.

McGinnis, C.S., Murrow, L.M., and Gartner, Z.J. (2019). DoubletFinder: Doublet Detection in Single-Cell RNA Sequencing Data Using Artificial Nearest Neighbors. Cell Syst 8, 329–337.e324.

Mesler, A.L., Benedeck, R.E., and Wong, S.Y. (2020). Preparing the hair follicle canal for hair shaft emergence. Exp Dermatol, doi: 10.1111/exd.14210.

Murphy, G.F., Lavker, R.M., Whitaker, D., and Korngold, R. (1991). Cytotoxic folliculitis in GvHD. Evidence of follicular stem cell injury and recovery. J Cutan Pathol 18, 309–314.

Niemann, C., and Horsley, V. (2012). Development and homeostasis of the sebaceous gland. Semin Cell Dev Biol 23, 928–936.

Page, M.E., Lombard, P., Ng, F., Gottgens, B., and Jensen, K.B. (2013). The epidermis comprises autonomous compartments maintained by distinct stem cell populations. Cell Stem Cell 13, 1–12.

Pan, X., Hobbs, R.P., and Coulombe, P.A. (2013). The expanding significance of keratin intermediate filaments in normal and diseased epithelia. Curr Opin Cell Biol 25, 47–56.

Panteleyev, A.A., Paus, R., Wanner, R., Nurnberg, W., Eichmuller, S., Thiel, R., Zhang, J., Henz, B.M., and Rosenbach, T. (1997). Keratin 17 gene expression during the murine hair cycle. J Invest Dermatol 108, 324–329.

Panteleyev, A.A., Rosenbach, T., Paus, R., and Christiano, A.M. (2000). The bulge is the source of cellular renewal in the sebaceous gland of mouse skin. Arch Dermatol Res 292, 573–576.

Parfitt, G.J., Lewis, P.N., Young, R.D., Richardson, A., Lyons, J.G., Girolamo, N.D., and Jester, J.V. (2016). Renewal of the Holocrine Meibomian Glands by Label-Retaining, Unipotent Epithelial Progenitors. Stem Cell Reports 7, 399–410.

Petersson, M., Brylka, H., Kraus, A., John, S., Rappl, G., Schettina, P., and Niemann, C. (2011). TCF/Lef1 activity controls establishment of diverse stem and progenitor cell compartments in mouse epidermis. EMBO J 30, 3004–3018.

Plewig, G., and Christophers, E. (1974). Renewal rate of human sebaceous glands. Acta Derm Venereol 54, 177–182.

Plewig, G., and Kligman, A.M. (1978). Proliferative activity of the sebaceous glands of the aged. J Invest Dermatol 70, 314–317.

Powell, A.E., Wang, Y., Li, Y., Poulin, E.J., Means, A.L., Washington, M.K., Higginbotham, J.N., Juchheim, A., Prasad, N., Levy, S.E., et al. (2012). The pan-ErbB negative regulator Lrig1 is an intestinal stem cell marker that functions as a tumor suppressor. Cell 149, 146–158.

Qiu, X., Hill, A., Packer, J., Lin, D., Ma, Y.A., and Trapnell, C. (2017a). Single-cell mRNA quantification and differential analysis with Census. Nat Methods 14, 309–315.

Qiu, X., Mao, Q., Tang, Y., Wang, L., Chawla, R., Pliner, H.A., and Trapnell, C. (2017b). Reversed graph embedding resolves complex single-cell trajectories. Nat Methods 14, 979–982.

Ramot, Y., Mastrofrancesco, A., Camera, E., Desreumaux, P., Paus, R., and Picardo, M. (2015). The role of PPARγ-mediated signalling in skin biology and pathology: new targets and opportunities for clinical dermatology. Exp Dermatol 24, 245–251.

Raymond, K., Richter, A., Kreft, M., Frijns, E., Janssen, H., Slijper, M., Praetzel-Wunder, S., Langbein, L., and Sonnenberg, A. (2010). Expression of the orphan protein Plet-1 during trichilemmal differentiation of anagen hair follicles. J Invest Dermatol 130, 1500–1513.

Reichenbach, B., Classon, J., Aida, T., Tanaka, K., Genander, M., and Göritz, C. (2018). Glutamate transporter Slc1a3 mediates inter-niche stem cell activation during skin growth. EMBO J 37, e98280.

Rittié, L., Tejasvi, T., Harms, P.W., Xing, X., Nair, R.P., Gudjonsson, J.E., Swindell, W.R., and Elder, J.T. (2016). Sebaceous Gland Atrophy in Psoriasis: An Explanation for Psoriatic Alopecia? J Invest Dermatol 136, 1792–1800.

Sardella, C., Winkler, C., Quignodon, L., Hardman, J.A., Toffoli, B., Attianese, G.M.P.G., Hundt, J.E., Michalik, L., Vinson, C.R., Paus, R., et al. (2018). Delayed hair follicle morphogenesis and hair follicle dystrophy in a lipoatrophy mouse model of Pparg total deletion. J Invest Dermatol 138, 500–510.

Schepeler, T., Page, M.E., and Jensen, K.B. (2014). Heterogeneity and plasticity of epidermal stem cells. Development 141, 2559–2567.

Schneider, M.R., and Paus, R. (2014). Deciphering the functions of the hair follicle infundibulum in skin physiology and disease. Cell Tissue Res 358, 697–704.

Selleri, S., Seltmann, H., Gariboldi, S., Shirai, Y.F., Balsari, A., Zouboulis, C.C., and Rumio, C. (2006). Doxorubicin-induced alopecia is associated with sebaceous gland degeneration. J Invest Dermatol 126, 711–720.

Shi, V.Y., Leo, M., Hassoun, L., Chahal, D.S., Maibach, H.I., and Sivamani, R.K. (2015). Role of sebaceous glands in inflammatory dermatoses. J Am Acad Dermatol 73, 856–863.

Singh, K., Camera, E., Krug, L., Basu, A., Pandey, R.K., Munir, S., Wlaschek, M., Kochanek, S., Schorpp-Kistner, M., Picardo, M., et al. (2018). JunB defines functional and structural integrity of the epidermo-pilosebaceous unit in the skin. Nat Commun 9, 3425.

Smith, K.R., and Thiboutot, D.M. (2008). Thematic review series: skin lipids. Sebaceous gland lipids: friend or foe? J Lipid Res 49, 271–281.

Stenn, K.S., Sundberg, J.P., and Sperling, L.C. (1999). Hair follicle biology, the sebaceous gland, and scarring alopecias. Arch Dermatol 135, 973–974.

Sundberg, J.P., Boggess, D., Sundberg, B.A., Eilertsen, K., Parimoo, S., Filippi, M., and Stenn, K. (2000). Asebia-2J (Scd1(ab2J)): a new allele and a model for scarring alopecia. Am J Pathol 156, 2067–2075.

Taetzsch, T., Brayman, V.L., and Valdez, G. (2018). FGF binding proteins (FGFBPs): Modulators of FGF signaling in the developing, adult, and stressed nervous system. Biochim Biophys Acta Mol Basis Dis 1864, 2983–2991.

Tang, W., Zeve, D., Suh, J.M., Bosnakovski, D., Kyba, M., Hammer, R.E., Tallquist, M.D., and Graff, J.M. (2008). White fat progenitor cells reside in the adipose vasculature. Science 322, 583–586.

Tassi, E., Garman, K.A., Schmidt, M.O., Ma, X., Kabbara, K.W., Uren, A., Tomita, Y., Goetz, R., Mohammadi, M., Wilcox, C.S., et al. (2018). Fibroblast Growth Factor Binding Protein 3 (FGFBP3) impacts carbohydrate and lipid metabolism. Sci Rep 8, 15973.

Vagnozzi, A.N., Reiter, J.F., and Wong, S.Y. (2015). Hair follicle and interfollicular epidermal stem cells make varying contributions to wound regeneration. Cell Cycle 14, 3408–3417.

Veniaminova, N.A., Grachtchouk, M., Doane, O.J., Peterson, J.K., Quigley, D.A., Lull, M.V., Pyrozhenko, D.V., Nair, R.R., Patrick, M.T., Balmain, A., et al. (2019). Niche-specific factors dynamically regulate sebaceous gland stem cells in the skin. Dev Cell 51, 326–340.

Veniaminova, N.A., Vagnozzi, A.N., Kopinke, D., Do, T.T., Murtaugh, L.C., Maillard, I., Dlugosz, A.A., Reiter, J.F., and Wong, S.Y. (2013). Keratin 79 identifies a novel population of migratory epithelial cells that initiates hair canal morphogenesis and regeneration. Development 140, 4870–4880.

Wang, S., Drummond, M.L., Guerrero-Juarez, C.F., Tarapore, E., MacLean, A.L., Stabell, A.R., Wu, S.C., Gutierrez, G., That, B.T., Benavente, C.A., et al. (2020). Single cell transcriptomics of human epidermis identifies basal stem cell transition states. Nat Commun 11, 4239.

Weinstein, G.D. (1974). Cell kinetics of human sebaceous glands. J Invest Dermatol 62, 144–146.

Yang, X., Zhong, X., Huang, A.J., and Reneker, L.W. (2021). Spontaneous acinar and ductal regrowth after meibomian gland atrophy induced by deletion of FGFR2 in a mouse model. Ocul Surf, In Press.

Yu, G., Wang, L.G., Han, Y., and He, Q.Y. (2012). clusterProfiler: an R package for comparing biological themes among gene clusters. OMICS 16, 284–287.

Zeeuwen, P.L., van Vlijmen-Willems, I.M., Hendriks, W., Merkx, G.F., and Schalkwijk, J. (2002). A null mutation in the cystatin M/E gene of ichq mice causes juvenile lethality and defects in epidermal cornification. Hum Mol Genet 11, 2867–2875.

Zhang, C., Chinnappan, M., Prestwood, C.A., Edwards, M., Artami, M., Thompson, B.M., Eckert, K.M., Vale, G., Zouboulis, C.C., McDonald, J.G., et al. (2021). Interleukins 4 and 13 drive lipid abnormalities in skin cells through regulation of sex steroid hormone synthesis. Proc Natl Acad Sci USA 118, e2100749118.

Zheng, Y., Eilertsen, K.J., Ge, L., Zhang, L., Sundberg, J.P., Prouty, S.M., Stenn, K.S., and Parimoo, S. (1999). Scd1 is expressed in sebaceous glands and is disrupted in the asebia mouse. Nat Genet 23, 268–270.

Zouboulis, C.C., and Boschnakow, A. (2001). Chronological ageing and photoageing of the human sebaceous gland. Clin Exp Dermatol 26, 600–607.

Zouboulis, C.C., Coenye, T., He, L., Kabashima, K., Kobayashi, T., Niemann, C., Nomura, T., Oláh, A., Picardo, M., Sasano, H., et al. (2022). Sebaceous immunobiology - skin homeostasis, pathophysiology, coordination of innate immunity and inflammatory response and disease associations. Front Immunol 13, 1029818.

