## Supplemental Figures S1-S7 for "Distinct mechanisms for sebaceous gland self-renewal and regeneration provide durability in response to injury"

### Veniaminova, Jia_Fig S1

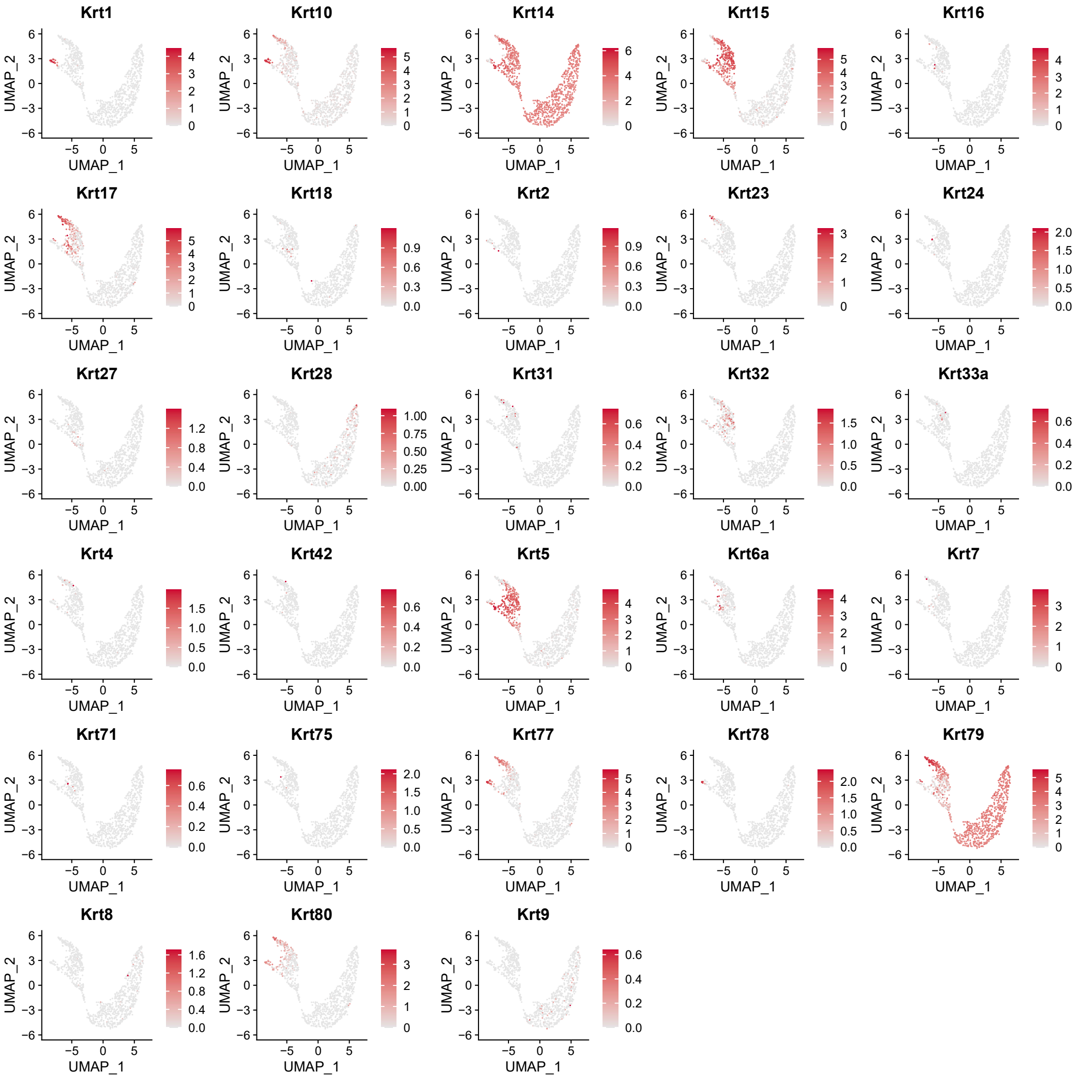

### Veniaminova, Jia_Fig S2

A

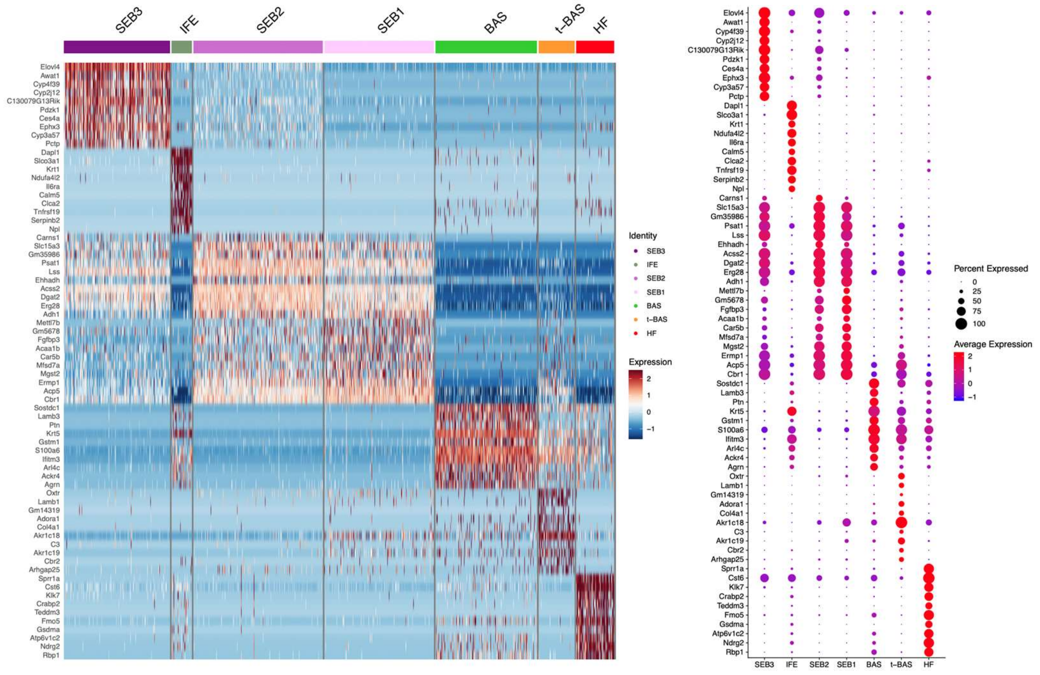


B

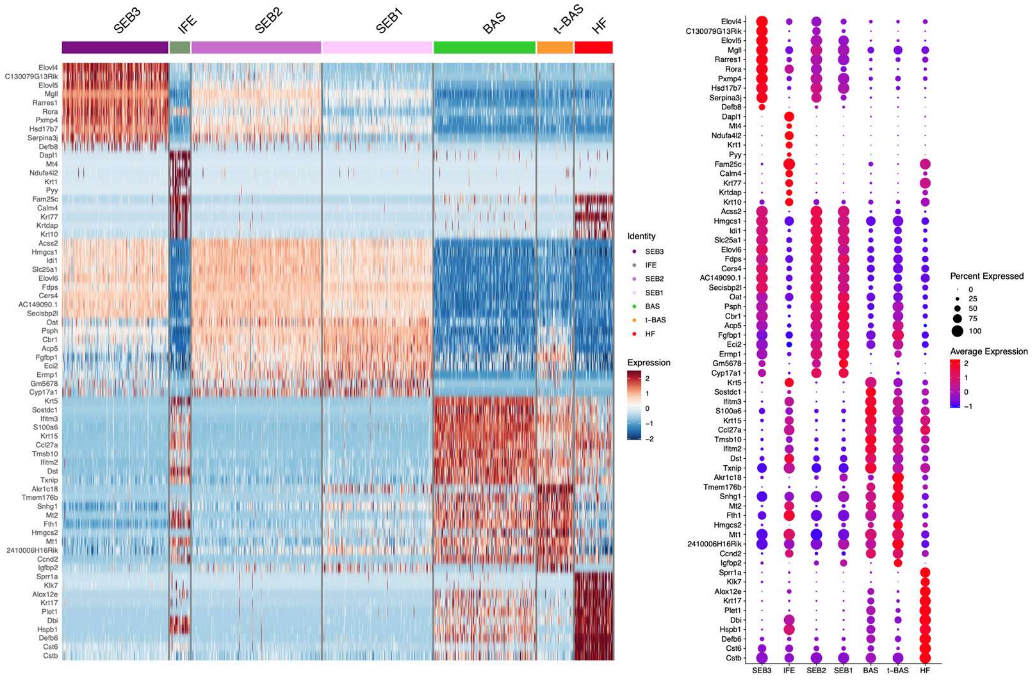

### Veniaminova, Jia_Fig S3

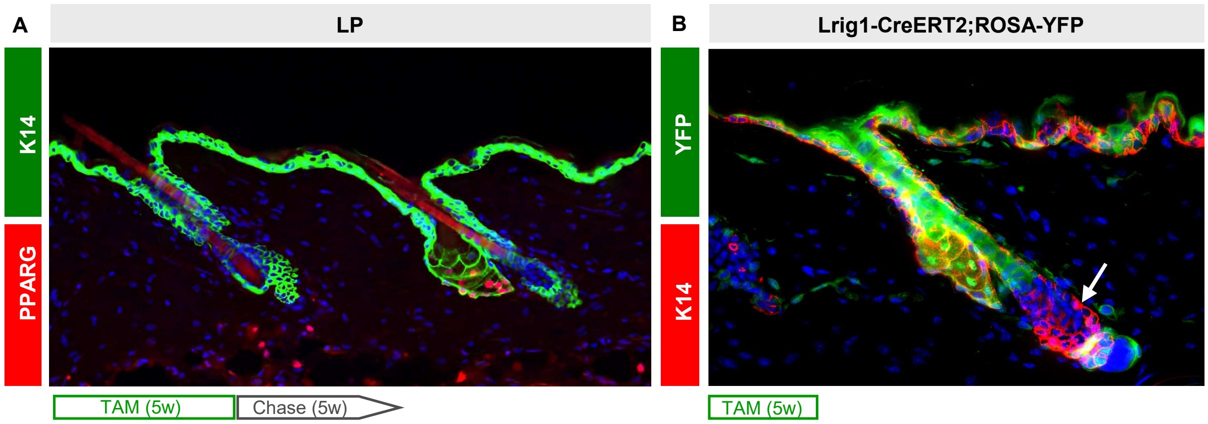

### Veniaminova, Jia_Fig S4

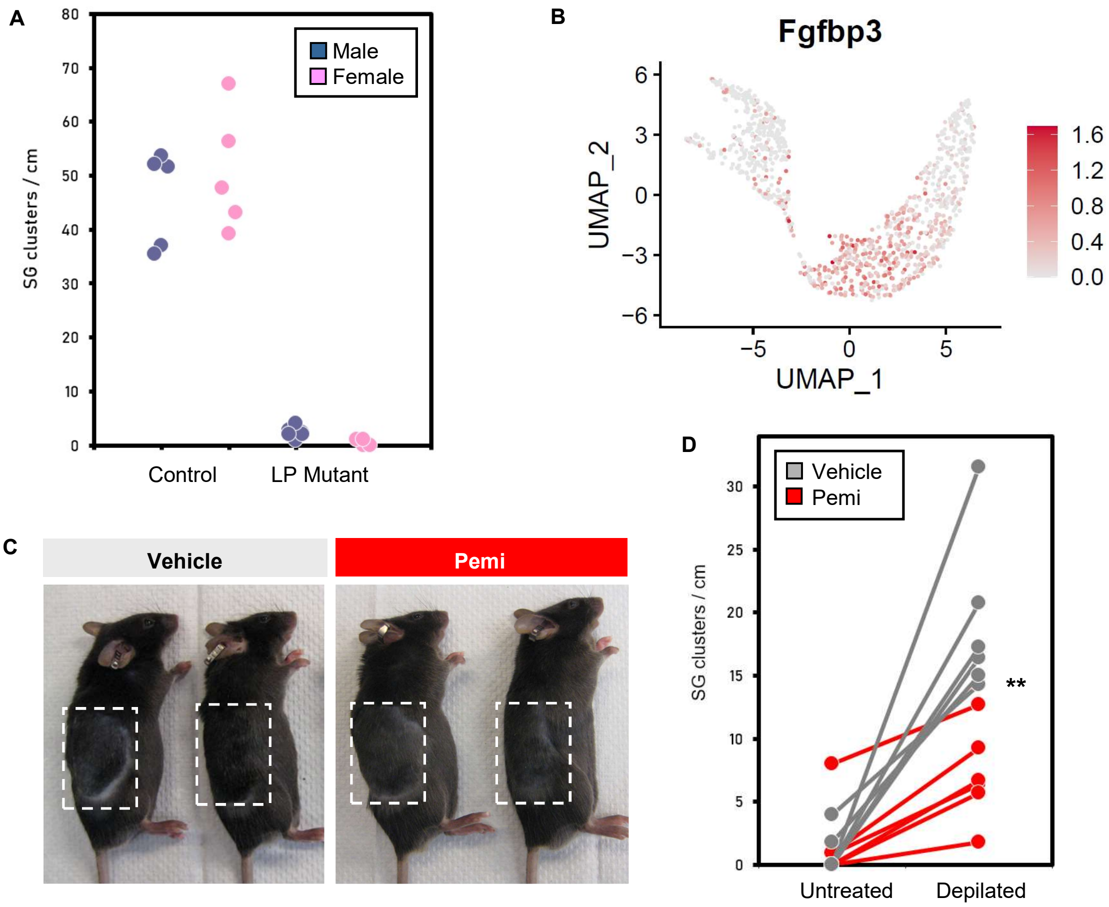

### Veniaminova, Jia_Fig S5

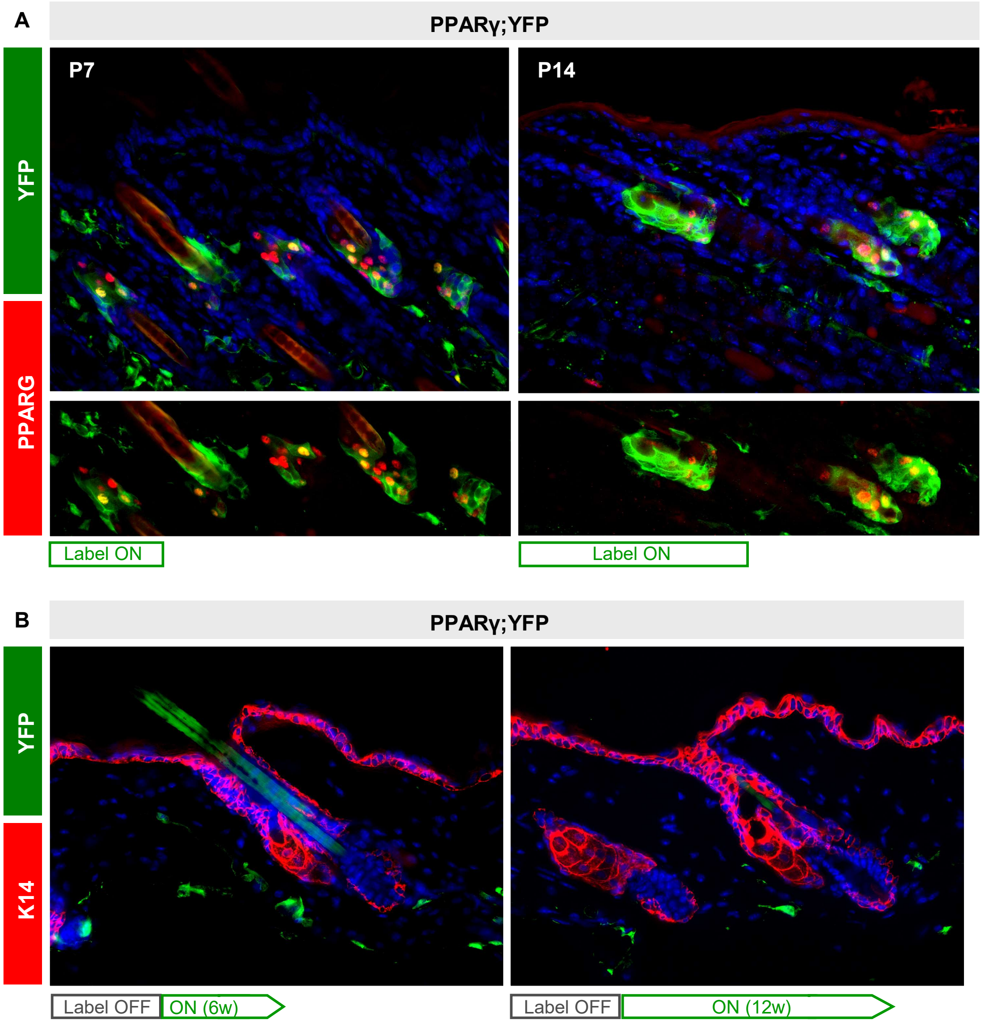

### Veniaminova, Jia_Fig S6

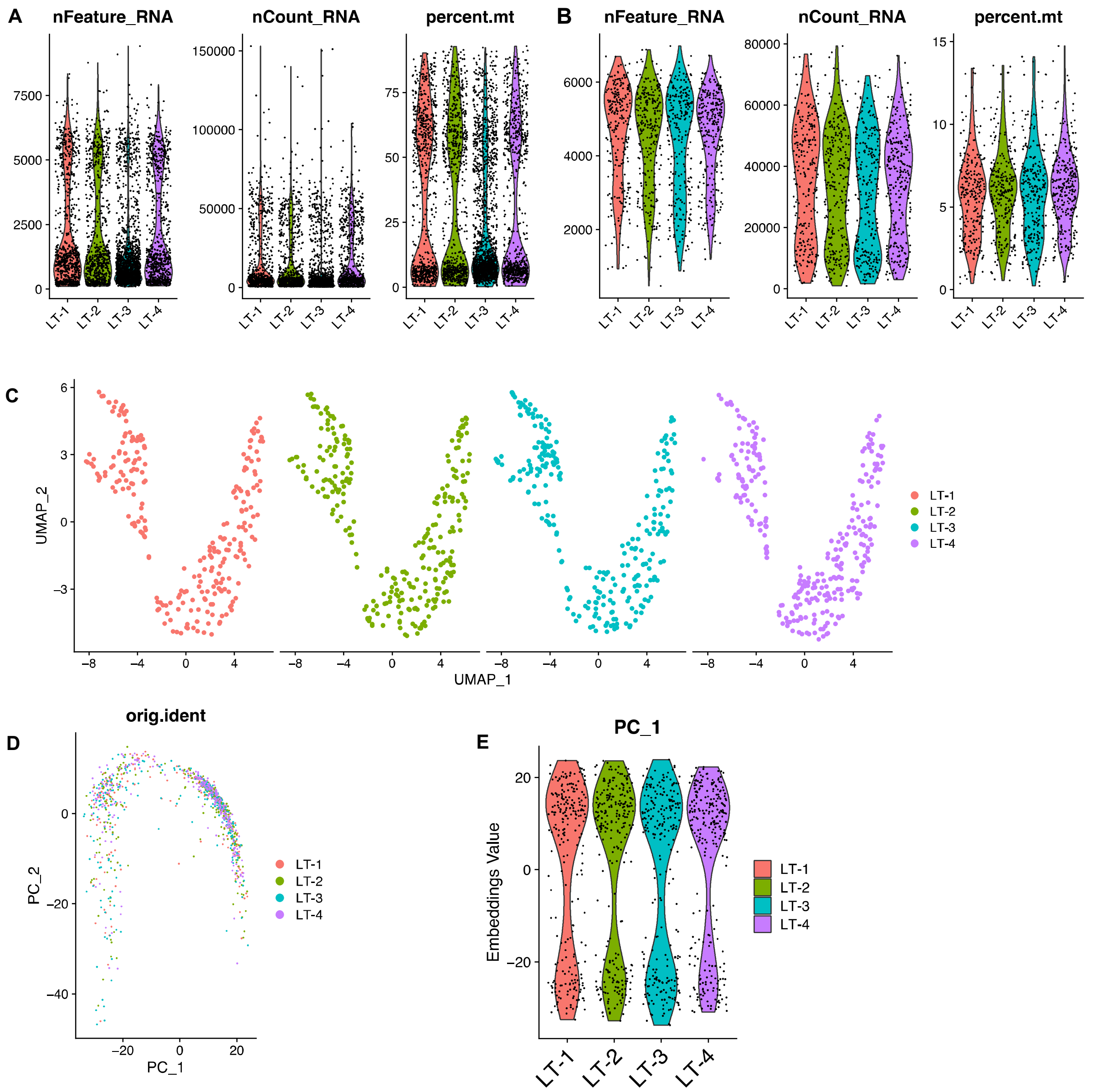

### Veniaminova, Jia_Fig S7

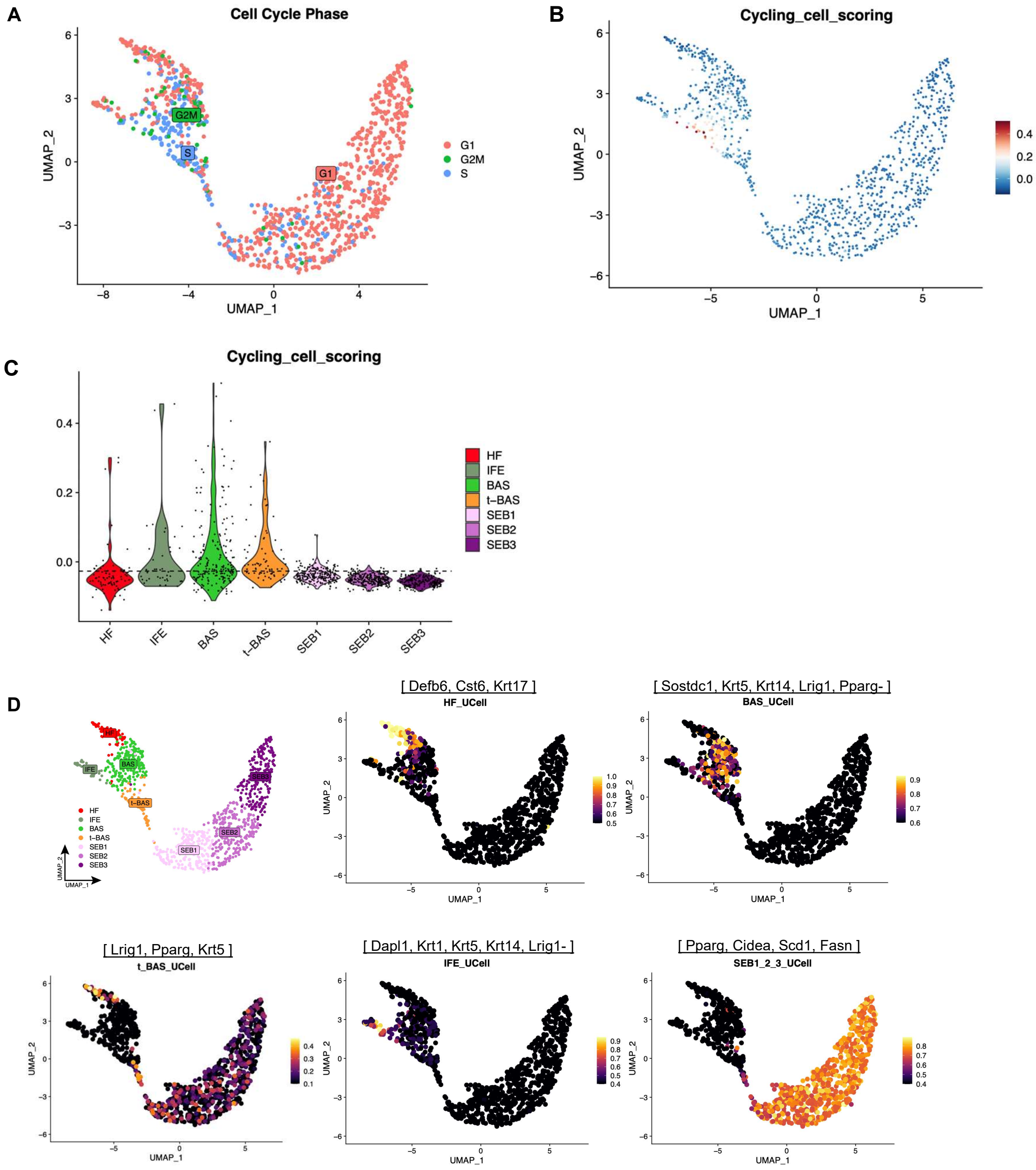
